# Magnetically Responsive Nanocultures for Direct Microbial Assessment in Soil Environments

**DOI:** 10.1101/2025.05.17.654660

**Authors:** Huda Usman, Mehdi Molaei, Stephen House, Martin F. Haase, Cindi L. Dennis, Tagbo H.R. Niepa

**Author notes:** These opinions, recommendations, findings, and conclusions do not necessarily reflect the views or policies of NIST, Sandia National Laboratory, or the United States Government. Certain equipment, instruments, software, or materials are identified in this paper in order to specify the experimental procedure adequately. Such identification is not intended to imply recommendation or endorsement of any product or service by the National Institute of Standards and Technology (NIST), nor is it intended to imply that the materials or equipment identified are necessarily the best available for the purpose.

## Abstract

Cultivating microorganisms in native-like conditions is vital for bioprospecting and accessing currently unculturable species. However, there remains a gap in scalable tools that can both mimic native microenvironments and enable targeted recovery of microbes from complex settings. Such approaches are essential to advance our understanding of microbial ecology, predict community functions, and discover novel biotherapeutics. We present magnetic nanocultures—a high-throughput microsystem for isolating and growing environmental microbes under near-native conditions. These nanoliter-scale bioreactors are encapsulated in semi-permeable membranes that form magnetic polymeric microcapsules using iron oxide nanoparticles within polydimethylsiloxane-based shells. This design offers mechanical stability and magnetic actuation, enabling efficient retrieval from soil-like environments. The nanocultures are optimized for optical and biological properties to support microbial encapsulation, growth, and sorting. Our study demonstrates the feasibility of using magnetically responsive microenvironments to cultivate elusive microbes, offering a promising platform for discovering previously uncultured or unknown microbial species.

**Teaser:** *Engineered magnetic nanocultures support microbial growth and magnetic separation from complex environments.*

## INTRODUCTION

Microorganisms are ubiquitous and play crucial roles in various ecosystems, such as nutrient cycling in soil and oxygen production in marine environments (*1–3*). Despite their known abundance, over 98 % of microbes remain undiscovered because they cannot be cultured using conventional laboratory methods (*4*). This phenomenon, known as “The Great Plate Count Anomaly,” highlights the significant gap between the number of microbes that can be observed microscopically and those that can be grown on agar plates (*5, 6*). Understanding these viable but nonculturable microorganisms is essential for uncovering new antibiotics, beneficial metabolites, and other therapeutic products (*7, 8*).

To address this challenge, microbial cultivation systems designed to mimic native environments are emerging as transformative tools in microbiology. These systems offer the potential to revolutionize sustainable agriculture, environmental conservation, and medical biotechnology by enabling the study of microorganisms under conditions closely resembling their natural habitats. However, replicating the complexity of natural ecosystems in controlled settings remains a significant challenge. Microbial communities interact with diverse environmental factors, and the lack of advanced tools to monitor and manipulate these conditions limits the effectiveness of current approaches (*9–15*).

Recent innovations, such as the isolation chip (iChip) and droplet-based microfluidics, have made strides in addressing these challenges. The iChip, for example, has facilitated the cultivation of up to 200 microbial species simultaneously by isolating single cells within micro-chambers separated by a semi-permeable membrane and incubating them directly in their native soil environments (*16–18*). This approach has provided groundbreaking insights into hard-to-cultivate species. However, iChips are limited by their high cost, micromachining complexity, and the need for physical placement in soil, which may restrict the range of microbial diversity that is captured (*19, 20*). Similarly, droplet microfluidics enables high-throughput isolation of single cells within water-in-oil (W/O) single emulsions, making it ideal for single-cell studies (*19, 21*). Despite these advantages, droplet systems often lack the physical stability and semi-permeability required for long-term studies and precise environmental control. Specifically, they are susceptible to coalescence and structural breakdown over time, which limits their use in extended incubations or complex environments (*20, 22*).

To overcome these limitations, we drew inspiration from both iChip and microfluidics to develop a versatile microbial cultivation platform capable of harbouring microbes in soil environments. In this study, we introduce magnetic nanocultures (MNCs), a novel platform designed to combine the high-throughput, customizable nature of droplet microfluidics with the permeability and environmental interaction capabilities of iChip. The magnetic nanoculture approach offers a key advantage: It bypasses the need for microfabricated chamber arrays by spontaneously forming each nanoculture via high-throughput droplet generation.

Magnetic nanocultures are nanoliter-sized water-in-oil-in-water (W/O/W) double emulsions, where the oil phase is formulated to crosslink into semi-permeable polydimethylsiloxane (PDMS)-based shell and house microbial cells. Throughout this study, we refer to “nanocultures” when discussing double emulsions that encapsulate microbial cells, and “microcapsules” when referring to similar structures containing only sterile water in the core. By incorporating magnetic nanoparticles (MNPs) into the PDMS structure, these nanocultures become mobilizable, allowing precise placement and retrieval from complex environments such as soil or marine ecosystems using magnetic actuation. This magnetic functionality is critical: without it, the nanocultures would be difficult to locate and recover after incubation periods, risking a loss of valuable samples. The magnetic nanoculture platform overcomes key limitations of conventional systems by incorporating a crosslinked, semi-permeable PDMS-based membrane that surrounds each aqueous core, providing structural integrity and enabling nutrient exchange, waste removal, and small-molecule diffusion—features essential for microbial viability and cross-communication (*9, 20*). Unlike traditional W/O emulsions, which are prone to coalescence and lack mechanical stability, or iChip platforms that require complex microfabrication and manual assembly of hundreds of chambers, magnetic nanocultures are spontaneously formed via high-throughput droplet generation in a capillary microfluidic device. Additionally, magnetic responsiveness enables targeted recovery from complex substrates such as soil—an advantage not readily achieved with existing methods. These technical improvements position this platform as a versatile and scalable tool for in situ microbial cultivation and retrieval, with broad potential in decoding environmental microbiomes, discovering previously unculturable species, and advancing microbial therapeutic development.

Building upon this foundation, we addressed critical challenges in microbial cultivation through several key advancements. We optimized the optical and magnetic properties of the nanocultures to ensure compatibility with imaging techniques and reliable magnetic actuation. Additionally, we preserved cell viability within the nanocultures, enabling robust growth and interaction studies under controlled conditions. We also achieved precise control over nanoculture movement using external magnetic fields, facilitating targeted placement and retrieval. Finally, we developed an efficient method for separating nanocultures from complex environments, such as soil samples, using their magnetic properties to streamline downstream analysis. Collectively, these advancements position the MNC platform as a versatile tool for *in situ* microbial cultivation, with transformative potential for microbiome research and biotechnological applications.

## RESULTS

### Characterization of Magnetic Nanoparticles and Functionalized Polymer Membranes

To functionalize the polymer with magnetic nanoparticles (MNPs), three nanoparticle diameters of 5 nm, 10 nm, and 20 nmwere evaluated for their suitability. Transmission electron microscopy (TEM) was used to assess the size distribution and uniformity of the nanoparticles (**Supporting Information, Fig. S1**). Representative high-angle annular dark-field scanning TEM images of the 5 nm, 10 nm, and 20 nm MNPs are shown in **Fig. S1 A**, **B**, and **C**, respectively. These images reveal distinct variations in size and morphology (**Fig. S1 D**), as further summarized in **Table S1**.

The 5 nm and 10 nm nanoparticles exhibited relatively uniform, spherical shapes with narrow size distributions of 4.7 +/- 1 nm and 9.3 +/- 0.9 nm, respectively. In contrast, the 20 nm nanoparticles displayed a broader size distribution of 14.8 +/- 6 nm and irregular morphologies, ranging from circular to faceted shapes (*p<0.001*, for all pairwise comparison amongst MNPs). These variations could influence the functionalization and performance of the polymer.

Additional characterization, including selected area electron diffraction patterns and energy-dispersive X-ray spectroscopy (**Supporting Information, Fig. S2**), confirmed that all three nanoparticle samples were crystalline iron oxide, which is magnetoresponsive (*23, 24*). This structural confirmation validates their suitability as candidates for imparting magnetic properties to the polymeric nanocultures.

Next, we sought to determine whether the MNPs could be uniformly incorporated into the polymer matrix to impart real-time manipulation, while maintaining optical transparency, which is essential for nanocultures, as it enables microscopic observation of microbial growth. Since we anticipated that the addition of MNPs could compromise optical transparency, we investigated how the incorporation of MNPs affected the optical properties of polymeric membranes (**Fig. 1**). To explore this, we generated various polymer blends using a 0.6:1 molar ratio of hydride (HMS-053) to vinyl (DMS-V21), with a platinum catalyst to initiate the hydrosilylation reaction in the presence of MNPs (**Fig. 1 A)**. The resulting PDMS blends containing 0 ppm (not shown), 60 ppm, 125 ppm, 250 ppm, and 500 ppm of MNPs were prepared and cured in standard cuvettes to assess their optical properties. Across the visible spectrum (385 nm–750 nm), transparency decreased with increasing MNP concentration and size (**Fig. 1 B1–B2)**. While pure PDMS transmitted 100 % of light (**Supporting Information, Fig. S3 A**), adding just 60 ppm of MNPs (regardless of size) reduced transmittance by 50 % to 23 % over the same range (*p < 0.0001*, compared to blank PDMS). At higher MNP concentrations, the visible light transmission was almost completely blocked for a path length of 1 cm in samples containing 20 nm, 10 nm, and 5 nm MNPs, respectively.

**Fig. 1.**
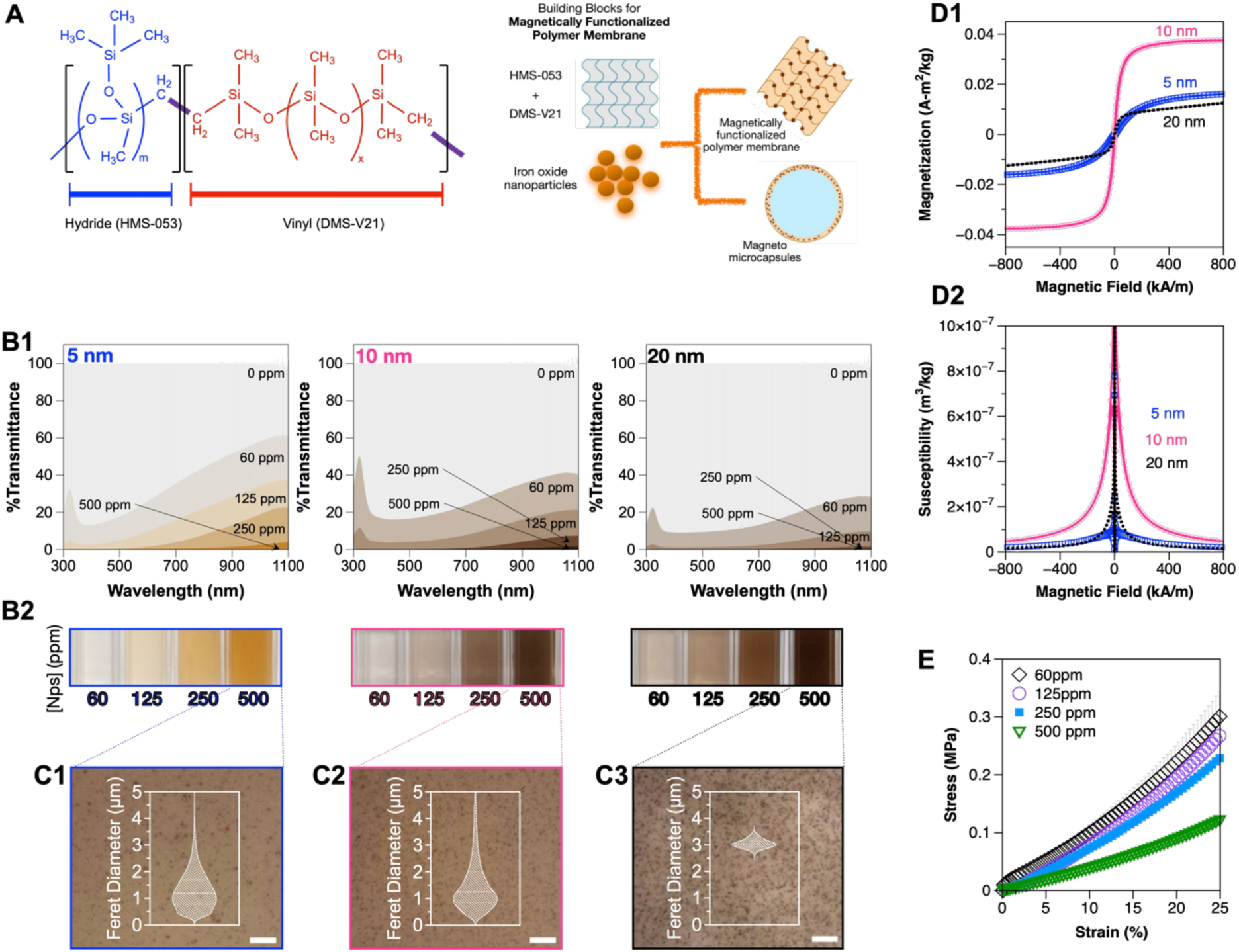
Characterization of PDMS Polymer Blends Containing Magnetic Nanoparticles (MNPs) **(A)** Schematic of the polymer crosslinking chemistry and the incorporation of MNCs into the PDMS matrix. (**B**) UV-Vis transmittance spectra (**B1**) and optical images (**B2**) of PDMS blends containing increasing concentrations (60 ppm, 125 ppm, 250 ppm, and 500 ppm) of 5 nm, 10 nm, and 20 nm magnetic nanoparticles (MNPs), showing a concentration-dependent decrease in transmittance and a corresponding increase in visual color intensity. Feret diameter distribution of MNP aggregates at 500 ppm, comparing 5 nm (**C3**), 10 nm (**C2**), and 20 nm (**C3**) sizes, demonstrating increasing aggregate size with nanoparticle size (scale bar: 20 µm). (**D1**) Magnetization and susceptibility (**D2**) curves of PDMS blends containing 500 ppm of each MNP size, demonstrating magnetic responsiveness of the composites. (**E**) Rheological analysis of PDMS blends containing 5 nm MNPs at varying concentrations, showing flow behavior as a function of shear rate. Standard deviations are shown, and the difference across the samples are statistically significant.

Given that the shell thickness of the nanocultures is approximately 9.8 µm +/- 2.2 µm, as determined from scanning electron microscopy (SEM) images of sectioned nanocultures (see Figure 2), we hypothesized that a path length shorter than 1 cm would increase transmittance, as predicted by Beer’s law. To test this, we used a UV-Vis spectrophotometer, this time rotating the cuvette by 90° to achieve a path length of 4 mm. As expected, reducing the path length resulted in a 20 % to 40% increase in percent transmittance for all MNP concentrations (**Supporting Information, Fig. S3 B**). These findings confirm that while MNP incorporation compromises optical transparency in bulk membranes, thinner layers, such as those used in nanocultures, retain sufficient transparency for microbial observation using both light and fluorescence microscopy.

**Fig. 2.**
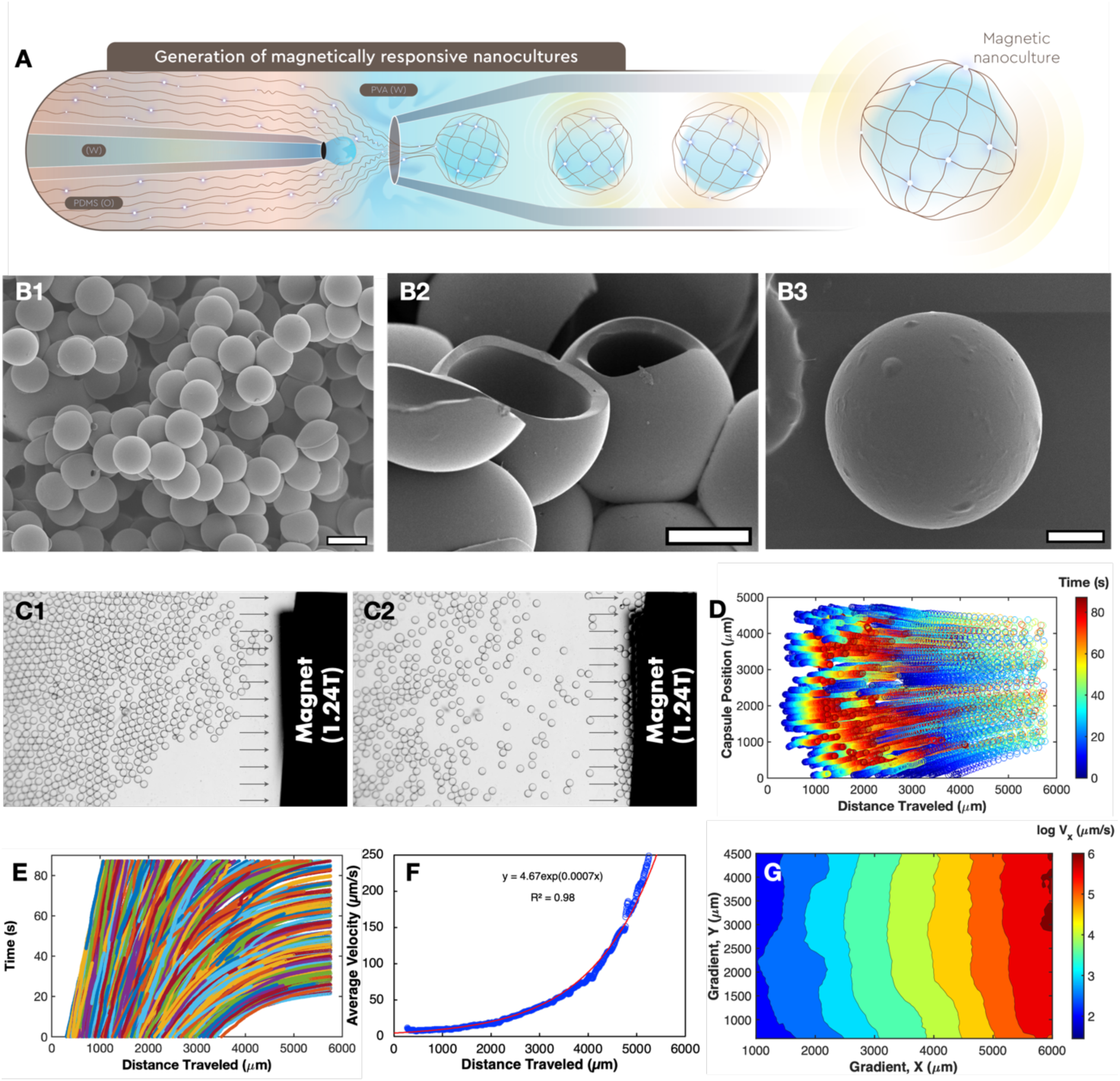
Generation and characterization of magnetically responsive microcapsules. (**A**) Generation of magnetically responsive nanocultures. The 3 liquid phases (aqueous culture medium, actuated PDMS mixtures, and PVA solution) were introduced into the microfluidic device and directed into a 3-phase interface for high-throughput generation of W/O/W emulsions. SEM images of nanocultures without (**B1-B2**) and with (**B3**) 500 ppm 5 nm MNPs. Scale bar: 200 μm, 100 μm, and 50 μm respectively, (**C1-C2**) MNCs undergoing magnetophoresis in the presence of external magnetic field. Scale bar: 500 µm (**D**) Magnetophoretic characterization of 500 ppm 5nm MNCs. (**E**) MATLAB rendering of MNC position versus distance traveled at a specified time. The plot of distance traveled towards the magnet of individual MNCs as a function of time. (**F**) Plot of average velocity as a function of distance traveled towards the magnet. (**G**) Velocity field gradient established within the channel in X and Y directions. The color map indicates the value of the average velocity in the x-direction.

After evaluating the effect of MNP concentration and size on optical transmittance, we next investigated how the distribution and uniformity of MNPs within the polymer matrix varied across MNP sizes. To explore this, membranes approximately 500 µm thick containing 500 ppm of 5 nm, 10 nm, and 20 nm MNPs were prepared and imaged at 1000× magnification using a Keyence profilometer optical microscope (**Fig. 1 C1-C3**). Feret’s diameter for the MNP aggregates was analyzed using ImageJ, revealing sizes of 1.1 µm +/- 0.9 µm, 1 µm +/- 3 µm, and 3.1 µm +/- 0.2 µm for 5 nm, 10 nm, and 20 nm MNPs, respectively (**inset, Fig. 1 C1-C3**). These aggregate sizes correspond approximately to ∼8 million 5 nm particles, ∼1.4 million 10 nm particles, and ∼0.87 million 20 nm particles per aggregate, assuming 75 % packing efficiency. Given the large errors and broad distribution, it can be inferred that MNPs were poorly dispersed in all cases. Based on prior studies of nanoparticle aggregation in viscous polymer systems, aggregate size is known to be sensitive to processing conditions (*25, 26*), suggesting that transparency versus magnetic susceptibility could be further tuned in future work by optimizing formulation and processing parameters.

Following the evaluation of MNP distribution and their impact on transparency, we next assessed whether embedding the MNPs into the polymer matrix would hinder their magnetic functionality, a critical feature for microcapsule actuation. To quantify the magnetic response of the MNP-containing samples, we measured the magnetic moment as a function of the applied magnetic field. These measurements were performed at multiple temperatures (277 K, 310 K, and 323 K) to assess potential environmental effects; however, little variation in magnetization was observed across conditions **(Supporting Information, Fig. S4 A-C)**. Therefore, we report representative data collected at 37 °C for samples containing 500 ppm of 5 nm, 10 nm, and 20 nm MNPs (**Fig. 1 D1**), which reflects the temperature used in most of our experiments. In contrast, magnetization increased with increasing MNP concentration **(Supporting Information, Fig. S4 D-F).** The 10 nm MNPs exhibited the highest saturation magnetization, followed by the 5 nm and 20 nm MNPs. The measured coercivity (3.2 kA/m) slightly exceeded the flux trappage (2.8 kA/m) in the superconducting magnet, indicating that the MNPs were not (super-)paramagnetic. For further analysis, the magnetic susceptibility χ (H) is plotted in **Fig. 1D2** to evaluate how the nanoparticles respond to the applied magnetic field and to quantify the extent of their magnetic behavior within the polymer matrix. A higher susceptibility indicates a stronger magnetic response, essential for ensuring that the microcapsules can be effectively actuated by an external magnetic field. Even though 10 nm exhibited higher susceptibility to the magnetic field compared to 5 nm and 20 nm, the polymer membrane containing 500 ppm of 5 nm MNPs exhibited the highest optical transparency while generating small MNP aggregates, making it the most favorable for microbial encapsulation purposes.

Beyond optical and magnetic characterization, we evaluated the mechanical properties of the MNP-containing polymer blends to ensure that the nanocultures could withstand environmental stresses during *in situ* incubation and retrieval. Compression testing was used to assess how varying concentrations of MNPs affect the stiffness and flow behavior of the polymer matrix.

As shown in **Fig. 1E**, increasing 5 nm MNP concentration from 60 ppm to 250 ppm led to progressively higher stress responses at a given shear rate, indicating increased stiffness and reduced flowability. However, at 500 ppm, the stress values decreased relative to 250 ppm, suggesting a loss in reinforcement efficiency at high nanoparticle loading. This trend may be attributed to nanoparticle aggregation or interfacial slippage, which can disrupt stress transfer within the polymer matrix—an effect previously observed in other nanoparticle-polymer systems (*27, 28*). These findings highlight a key design consideration for functionalized polymer membranes: Magnetic nanoparticle loading must be carefully optimized to balance mechanical robustness with properties suitable for microbial cultivation and retrieval in soil environments. This trade-off prompts a broader question relevant to platform development:

*How can one achieve sufficient optical transparency for real-time observation while ensuring the mechanical integrity required to withstand environmental stresses?* Mechanical behavior, including stiffness and flow resistance, is critical for applications where the polymer must endure environmental stress while maintaining the optical clarity necessary for real-time monitoring. Our results suggest that while increasing MNP concentration enhances magnetic responsiveness and stiffness, it may also reduce flexibility and transparency, potentially limiting performance in applications that demand a careful balance between structural integrity and imaging compatibility.

Lastly, microbial encapsulation requires the containment of an aqueous suspension in hydrophobic polymer membranes; therefore, it was essential to prove that the addition of the MNPs did not alter the surface hydrophobicity of the polymer membranes (*20, 22*). To test this, actuated PDMS membranes were prepared by vortexing polymer mixtures with MNPs, followed by degassing and curing, as described in the Materials and Methods section. The contact angle was determined by analyzing the shape of the droplet about the three-phase contact line between the solid membrane, liquid droplet, and ambient air. In such measurement, hydrophilicity is indicated by a contact angle smaller than 90°, whereas hydrophobicity is indicated by a contact angle larger than 90°. The surface contact angle measurements of the actuated PDMS with a variable concentration of MNPs (from 60 ppm to 500 ppm) confirmed that, in all cases, the PDMS remains hydrophobic **(Supporting Information, Fig. S5,** *differences of p < 0.05 were considered statistically significant*). Specifically, the contact angles were 101° +/- 4°, 108° +/- 4°, and 106° +/- 3° for the 5 nm, 10 nm, and 20 nm MNPs, respectively **(Fig. S5 A, B, C)**. These measurements were comparable with the background PDMS (without any MNPs) samples, for which the contact angle was about 110°. The surface contact angle measurements confirmed that the hydrophobic nature of the PDMS membrane was preserved across different MNP concentrations. This ensures the membrane’s ability to contain aqueous microbial cultures effectively.

Based on the combined analysis of optical transparency, magnetic responsiveness, and mechanical properties, the polymer membrane containing 500 ppm of 5 nm MNPs was selected as the optimal formulation for further experimentation. While the 10 nm nanoparticles exhibited higher magnetic susceptibility, the 5 nm nanoparticles provided the best balance between optical transparency and mechanical strength. This made them particularly suitable for microbial encapsulation applications, where transparency is critical for real-time monitoring of microbial growth. Moving forward, all subsequent experiments involving the generation of microcapsules or nanocultures were conducted using 500 ppm of 5 nm nanoparticles, unless otherwise specified. In the following section, we explore the development of magnetically responsive microcapsules.

### Generation and Characterization of Magnetically Responsive Microcapsules

Based on the favorable optical, mechanical, and magnetic properties of the PDMS-MNP blends, we hypothesized that the polymer mixture actuated with 5 nm MNP could be used to generate stable W/O/W double emulsions and serve as the basis for magnetic microcapsules and nanocultures. To test this, we employed a co-flow focusing glass capillary microfluidic device to create monodisperse magnetic microcapsules, containing no bacteria (**Fig. 2**). The inner phase of magnetic microcapsules consisted of sterile water, the middle phase was PDMS containing 500 ppm of 5 nm MNPs, and the outer phase was 5 wt% PVA solution. The PDMS mixture, composed of hydride (HMS-053) and vinyl (DMS-V21) groups mixed at a 0.6:1 molar ratio and catalysed with 0.5 ppm Pt, was thermally crosslinked by heating at 70 °C for 5 min, followed by incubation at 37 °C for 24 h to mimic bacterial culturing conditions. The heat-pretreatment was previously shown to have no effect of cell viability (*20, 22*). After 24 h, fully crosslinked microcapsules were stored at room temperature for downstream experiments such as magnetophoresis.

These magnetically responsive microcapsules were then characterized for their morphology, size distribution, shell thickness, and behavior under a magnetic field, with water as the inner phase in place of bacteria (**Fig. 2**). This characterization helped confirm the integrity and functionality of the polymeric shell embedded with MNPs. The schematic in **Fig. 2 A** illustrates how the external magnetic field interacts with the microcapsules, enabling remote actuation and guiding downstream recovery processes. This microcapsule fabrication protocol was subsequently used for generating microbial nanocultures, with microorganims in the core. **Fig. 2 B1-B3** show scanning electron microscopy (SEM) images of the microcapsules at different magnifications. These images confirm the successful formation of spherical and monodisperse nanoliter-sized microcapsules, with the diameters ranging from 180 to 200 µm, corresponding to a volume of 3 to 4.2 nL, and an average shell thickness of 9.8µm +/- 2.2 µm. **Fig. 2 B1** shows the microcapsules in bulk, while **Fig. 2 B2** and **2 B3** provide more detailed views of individual capsules, highlighting the smooth surface and encapsulation integrity, both without (**Fig. 2 B1-B2)** and with MNPs (**Fig. 2 B3)**.

**Fig. 2 C1-C2** demonstrates the magnetic actuation of the microcapsules when exposed to an external magnetic field. In the absence of a magnetic field (**Fig. 2 C1**), the nanocultures are evenly distributed throughout the collection channel. However, once a magnetic field is applied (**Fig. 2 C2**), the microcapsules respond by clustering and aligning along the field gradient, confirming their magnetic responsiveness **(Supporting Video 1-2)**. This raises a question: *Can this precise alignment and movement be controlled in real time to enhance applications such as targeted delivery and environmental sensing?*

A time-resolved plot of the microcapsule positions under the influence of the magnetic field illustrates their displacement over time, providing insight into how far they can travel within a given time frame—an important parameter for designing retrieval strategies in real-environment applications (**Fig. 2 D)**. In **Fig. 2 E**, the trajectories of individual microcapsules are plotted as a function of distance traveled and time. Each colored line represents a single microcapsule, and the trend reveals a consistent increase in velocity over time, reflecting magnetic acceleration along the field gradient. **Fig. 2 F** shows the average velocity of the microcapsules as a function of the distance they traveled, confirming a nearly exponential increase in velocity as the distance decreases from the magnet. This behavior aligns with the expected magnetic actuation, where microcapsules accelerate as they move deeper into the magnetic field gradient.

Next, we present in **Fig. 2 G** the gradient of the magnetic field as a function of the distance traveled, further illustrating the strong correlation between the magnetic field strength and the velocity of the microcapsules. The color gradient represents the velocity field of the microcapsules in the x-direction, with warmer colors indicating higher velocities. This gradient information is essential for controlling the actuation and movement of the microcapsules in applications such as targeted delivery or real-time manipulation.

Having characterized the morphology, magnetic actuation, and velocity profiles of the hollow microcapsules, we established their stability and behavior under magnetic fields. These findings demonstrate the precise control and responsiveness of the microcapsules, instilling confidence in their suitability for *in situ* incubation and retrieval. However, while the physical and magnetic properties are promising, their biological compatibility remains critical for broader applicability. Specifically, it is essential to determine whether the nanoparticles embedded in the polymer matrix exhibit cytotoxic effects on microbial cells. In the following section, we evaluate the cytotoxicity of MNP-containing polymer membranes and nanocultures to assess their safety for microbial encapsulation.

### MNP-Containing PDMS Membranes Exhibit No Antimicrobial Properties

The antimicrobial properties of the newly synthesized MNP-containing membranes were evaluated, as they are designed to house microorganisms. To assess their biological inertness, *Staphylococcus aureus* (SH1000, Gram-positive) and *Pseudomonas aeruginosa* (PAO1, Gram-negative) were cultured in ultra-filtered tryptone yeast extract (UFTYE) medium in the presence of the MNPs at varying concentrations incorporated into the membranes. Our results demonstrated that cells could survive exposure up to 500 ppm MNPs, which corresponds to the concentration used to generate the MNP-containing membranes (**Fig. 3**). No significant change in the growth potential of either cell type was observed, compared to the untreated control across all three MNP sizes (5 nm, 10 nm, and 20 nm). Statistical analysis indicated no significant differences (*p > 0.05* for all conditions compared to the control), confirming that the MNPs, at these concentrations, did not negatively impact bacteria growth **(Supporting Information, Fig. S6)**.

**Fig. 3.**
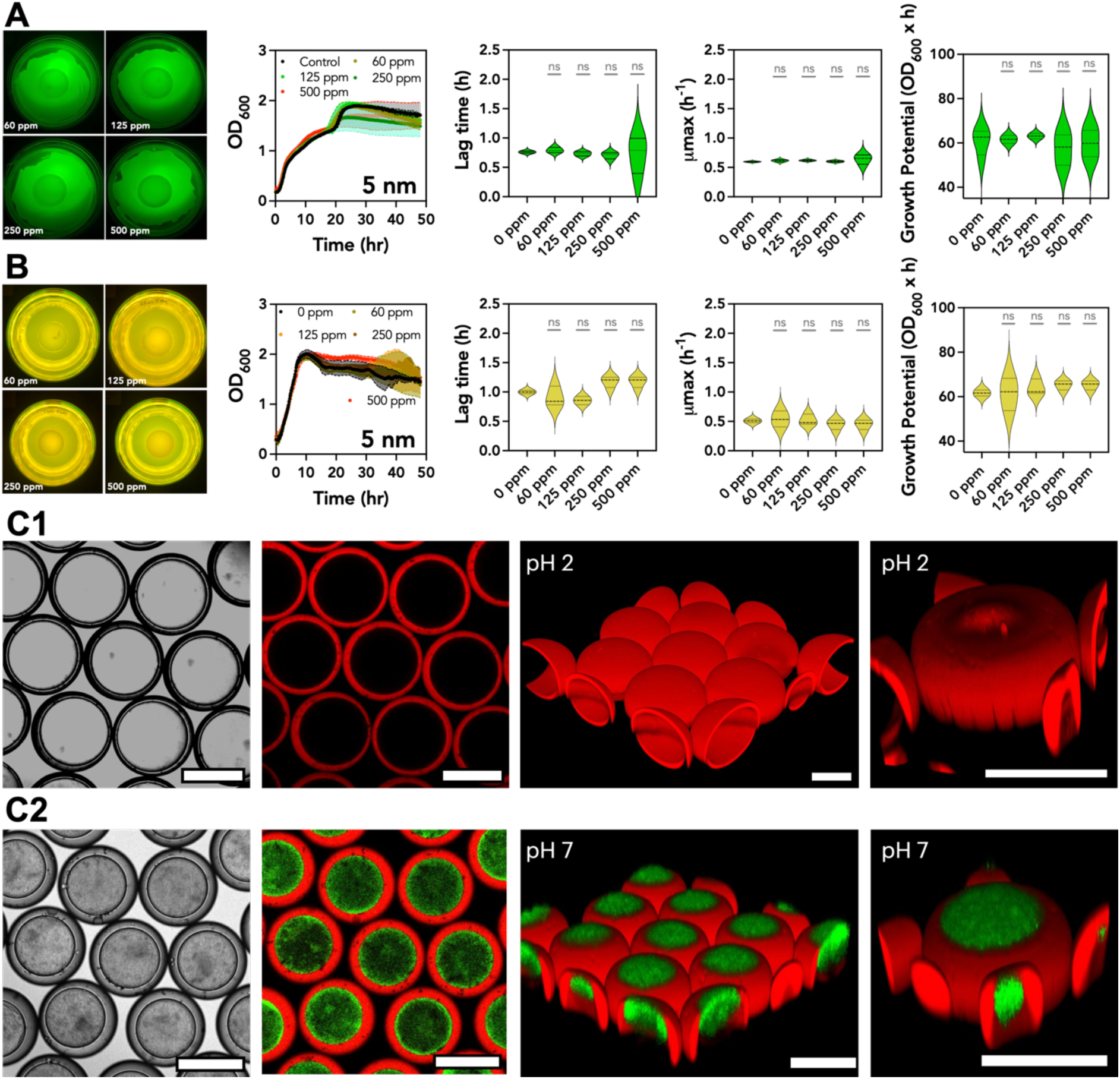
Cell susceptibility when exposed to varying concentrations of MNPs. (**A**) GFP-tagged *Staphylococcus aureus* and (**B**) *Pseudomonas aeruginosa* were exposed to MNP concentrations of 60 ppm, 125 ppm, 250 ppm, and 500 ppm, growth curves were constructed by taking optical density (OD) measurements every 10 min over 48 h at 600 nm. Differences were considered significant when p < 0.05, (**C**) GFP-tagged *P. aeruginosa* was encapsulated in the MNCs, and microbial growth was observed for 24 h before confocal imaging. The nanocultures were collected in **(C1)** pH = 2 and **(C2)** pH = 7 to assess the effects of their growth within the MNCs in simulated natural environments. Scale bar: 100 µm.

Similarly, when the MNP-containing membranes (with MNP concentrations ranging from 60 ppm to 500 ppm) were tested for their effect on cell viability, no significant impact was observed for either *S. aureus* or *P. aeruginosa* (**Fig. 3 A-B)**. Thin discs (10 mm in diameter and 2 mm thick) composed of the MNP-containing PDMS membranes were placed on Lysogeny Broth (LB) agar plates inoculated with SH1000 and PAO1. No inhibition zones were detected around the membranes, indicating the absence of antimicrobial activity. To further investigate the biological inertness of the membranes, Green Fluorescent Protein (GFP)-tagged PAO1 cells were encapsulated within the MNCs to determine whether the presence of MNPs in the shell would interfere with microbial growth. The shell of the nanoculture was stained with Nile Red for enhanced imaging. Confocal microscopy images in **Fig. 3 C1** and **3 C2** show the nanocultures collected at pH 2 and pH 7, representing simulated natural environments, such as under acid rain. These conditions enabled us to assess the effect of pH variation on microbial viability and growth within the nanocultures. At pH 2, microbial growth was significantly affected, as seen by the absence of green color in the confocal images, while at neutral pH (pH 7), the cells exhibited normal growth (confluent nanocultures), suggesting that the acidic environment, rather than the MNPs, was responsible for growth inhibition. Together, these findings confirm that the MNP-containing PDMS membranes exhibit sufficient optical transparency for real-time observation of fluorescently labelled microbes. They further demonstrate that the MNCs do not exhibit any antimicrobial properties, further supporting their biological inertness **(Supporting Information, Fig. S6-S8)**. This characteristic is crucial for applications where the membranes are used to encapsulate and support microbial growth without introducing cytotoxic effects.

### Microbial Growth Dynamics and Magnetic Actuation of Nanocultures

We further investigated the growth dynamics of *E. coli Nissle,* another Gram-negative microbial species, and the magnetic responsiveness of the resulting MNCs. We compared the velocity profiles of the bacteria-loaded MNCs to the empty microcapsules to assess whether bacterial encapsulation alters their movement under a magnetic field (**Fig. 4**). **Fig. 4 A** shows the growth dynamics of *E. coli Nissle* encapsulated within the MNCs over 24 h **(Supporting Video 3)**. Wide field images taken at different time intervals reveal that the encapsulated bacteria maintain their viability and proliferate within the MNCs. The gradual increase in cell density indicates healthy bacterial growth, suggesting that the nanoparticles embedded in the MNC shell do not inhibit bacterial viability, motility patterns, or replication. Additionally, the MNCs visibly shrink as bacterial growth progresses, suggesting that the membrane allows waste diffusion and supports a dynamic microenvironment conducive to microbial proliferation.

**Fig. 4:**
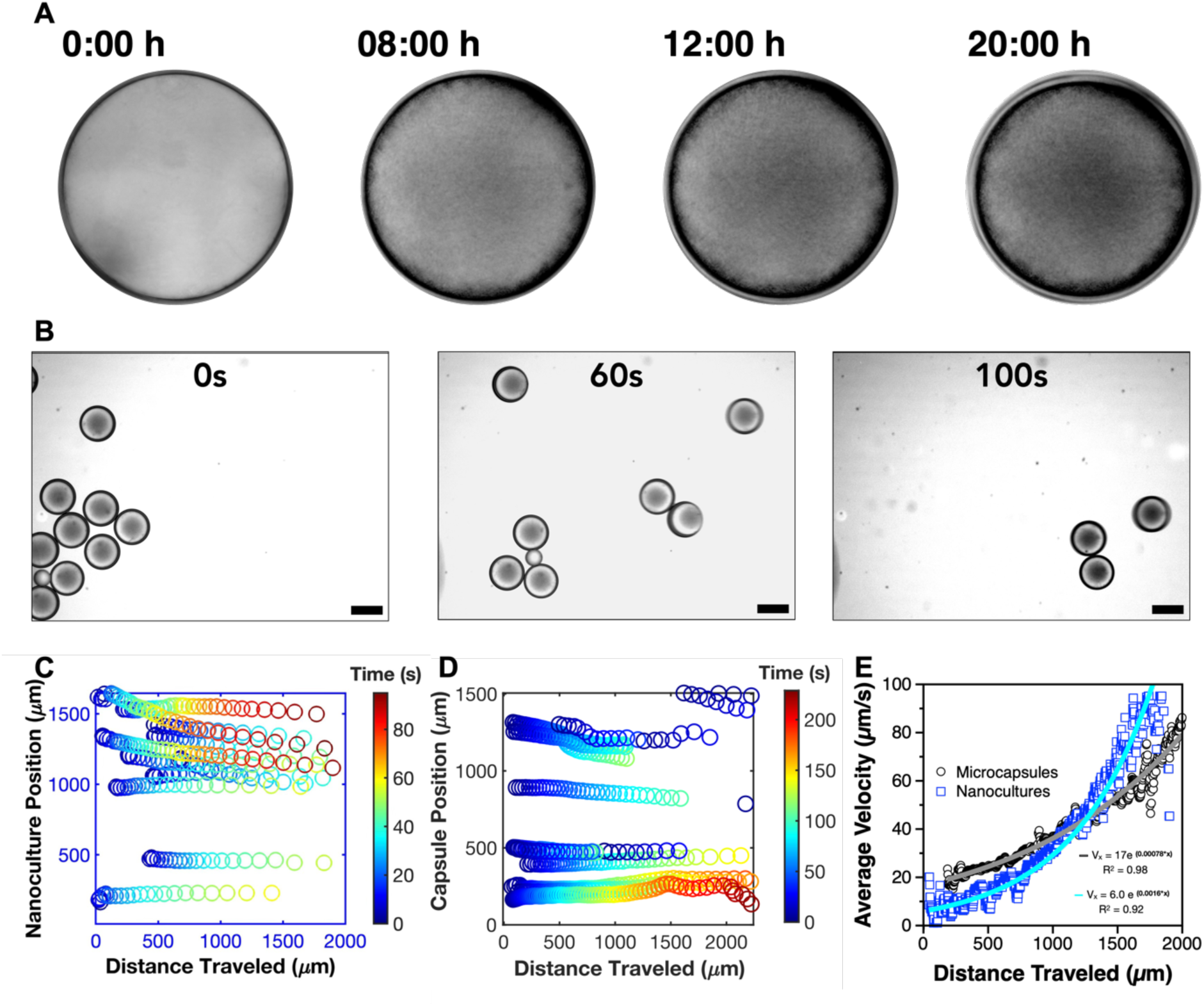
Microbial growth assessment and magnetic actuation of microcapsules. (**A**) (A) 24-hour growth assessment of wild-type *E. coli Nissle* in the MNCs. Scale bar: 50 µm. (**B**) Magnetic actuation of *E. coli Nissle* nanocultures, showing motion towards the magnet on the right. Scale bar: 200 μm. MATLAB rendering of the magnetic actuation of the (**C**) microcapsules vs. (**D**) MNCs showing faster motion in the later condition. (**E**) The average velocity gradients of the empty microcapsules vs the nanocultures is presented.

In **Fig. 4 B**, we examined the movement of MNCs containing *E. coli Nissle* under the influence of a magnetic field. The time-lapse images (taken at 0 s, 60 s, and 100 s) show that the MNCs initially start randomly distributed. As the magnetic field is applied, they rapidly move across the field of view in 100 s, and clustered near the magnetic source (**Fig. 4 C**). This movement highlights the strong magnetic properties of the MNC shell, which remain effective even in the presence of encapsulated bacteria **(Supporting Video 4)**.

The characterization of microcapsule velocity without encapsulated bacteria is shown in **Fig. 4 D** (**Supporting Video 5**), with the microcapsule positions presented as a function of the distance traveled over time. The time gradient reveals that the microcapsules move uniformly toward the magnetic source, but at a slower pace compared to the nanocultures. The comparison between the velocity gradient of the nanocultures and the microcapsules presented in **Fig. 4 E** shows that the presence of *E. coli Nissle* inside the MNCs does by no mean impede the actuation of the nanocultures. In both cases, the average velocity follows a similar exponential trend, with only slight variations in individual trajectories. This suggests that the added mass of the encapsulated bacteria is negligible relative to the overall mass of the MNCs and does not significantly alter magnetic actuation. In fact, cell metabolism could be aiding actuation if bacteria are secreting biosurfactants that lubricate the capsule surface and reduce drag along the wall or increase the apparent viscosity of the milieu (*29*). The ability of the MNCs to maintain consistent motion despite small payload differences highlights their robustness for applications such as microbial manipulation, targeted delivery, and environmental sensing.

### Broader Applications of Magnetically Responsive Nanocultures

Magnetic-based separation technologies have already found extensive applications across medical and industrial fields, including cancer cell detection (*30, 31*), apheresis (*32, 33*), and municipal water purification (*34, 35*). In this study, we developed an MNC system to address the growing need for high-throughput, low-cost, and straightforward microbial separation tools. After successfully cultivating bacteria within these MNCs, we extended the application to demonstrate the ability to magnetically sort different microbial species encapsulated in distinct artificial microenvironments (**Figs. 5 and 6**).

**Fig. 5:**
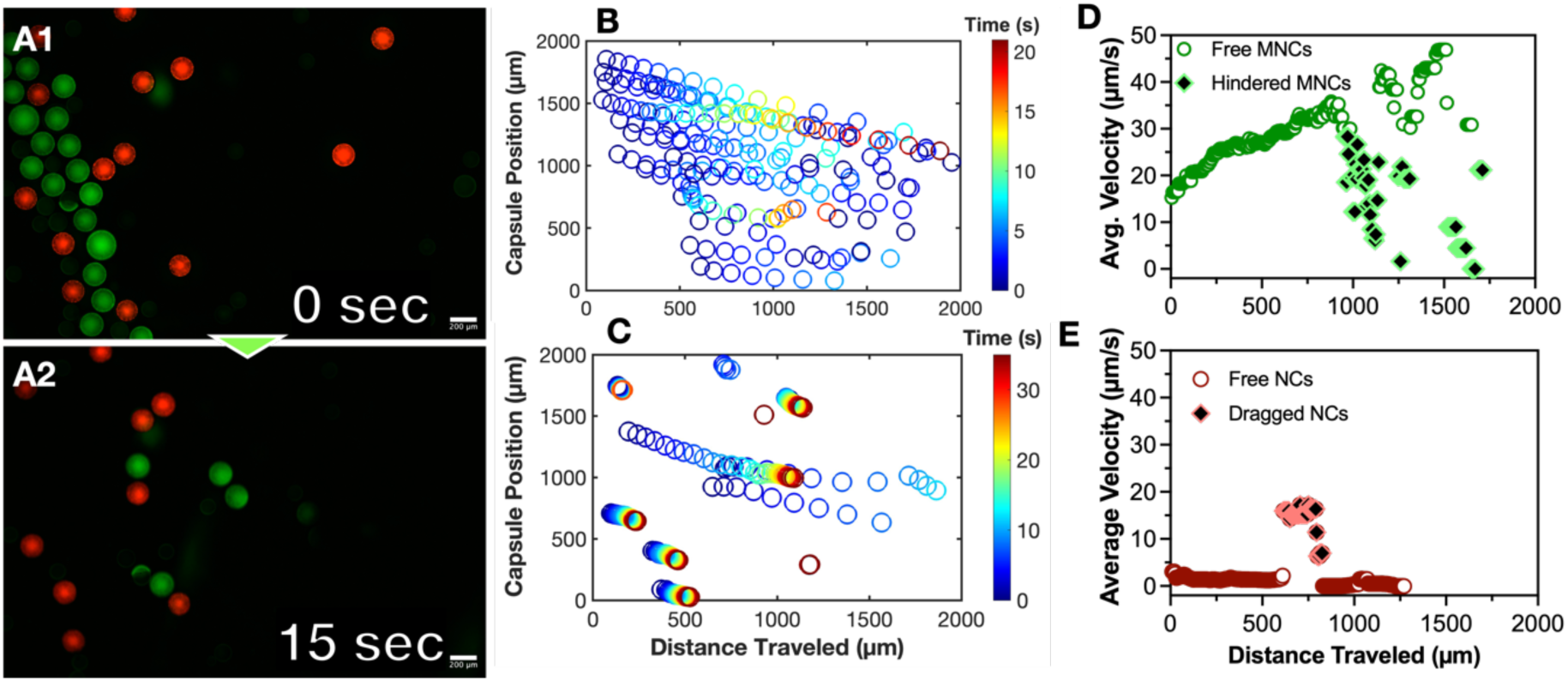
Magnetic sorting of nanocultures containing both MNP-actuated and non-actuated. (**A1-A2**) *E. coli Nissle* stained with Syto 9 (green fluorescence) and encapsulated in MNCs with 5 nm MNPs alongside RFP-tagged *E. coli Nissle* encapsulated in non-actuated nanocultures. Representative fluorescent microscopy images show the sorting process over time as the magnet attracts the MNP-actuated nanocultures. Scale bar: 200 μm. (**B-C**) MATLAB-rendered trajectories of actuated (green, **B**) and non-actuated (red, **C**) nanocultures as a function of distance traveled toward the magnet. (**D-E**) Average velocity values of the actuated (green, **D**) and non-actuated (red, **E**) nanocultures following actuation.

**Fig. 6:**
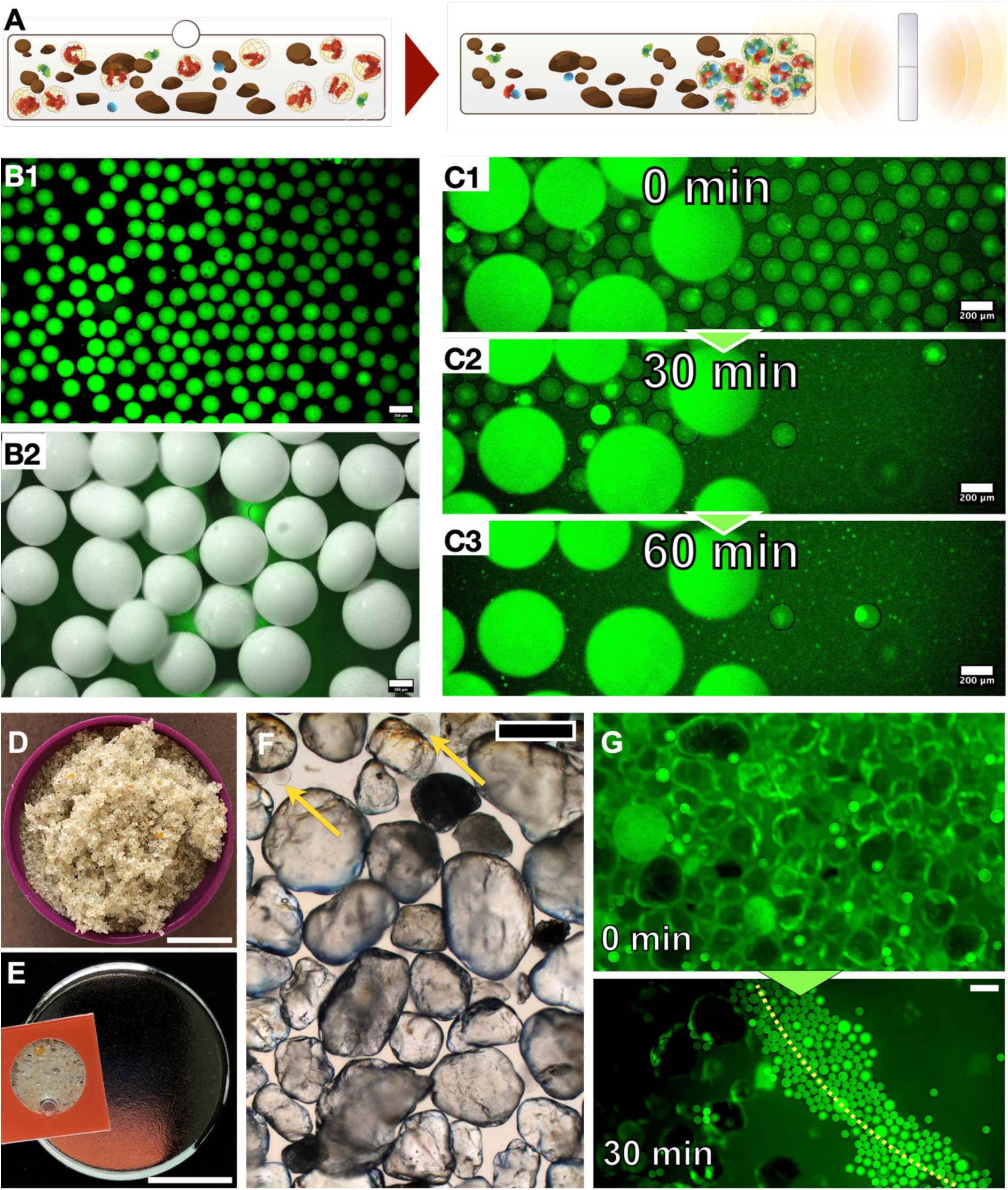
Real-world applicability of magnetic nanocultures. (**A**) Proof of concept for retrieving MNCs from an artificial environment. (**B1**) Image of *E. coli Nissle* encapsulated in MNCs stained with Syto 9. Scale bar: 200 µm. (**B2**) Image of 0.5 mm zirconia/silica beads introduced to replicate the spatial heterogeneity encountered in soil systems. Scale bar: 200 µm. (**C1-C3**) MNCs undergoing magnetophoresis. Magnet positioned to the right. Scale bar: 200 μm. (**D**) Beach sand was used for the experiments. Scale bar: 1 cm (**E**) Sand packed into the hybridization chamber for the retrieval of nanocultures. Scale bar: 0.5 in (**F**) Zoomed-in image of heterogenous sand particles. Scale bar: 200 µm (**G**) Nanocultures after retrieval from under the sand. Scale bar: 200 µm.

To test the efficacy of the MNC system in separating encapsulated microbes, we generated nanocultures containing *E. coli Nissle* functionalized with 500 ppm MNPs, while non-magnetic control nanocultures encapsulated Red Fluorescent Protein (RFP)-tagged *E. coli Nissle* (**Fig. 5 A)**. The MNCs containing *E. coli Nissle* were stained with Syto 9 to enhance visualization during magnetic separation (**Fig. 5 A1-A2)**. This dye was selected for its ability to partition through PDMS nanoculture shells and permeate bacterial membranes, staining the total nucleic acid green (*9, 21*). We hypothesized that the MNCs would be efficiently separated from their non-magnetic counterparts. To test this, both the functionalized and non-functionalized nanocultures were placed in a sealed chamber filled with ultrafiltered tryptone-yeast extract (UFTYE) medium, and a neodymium magnet with a pull strength of 0.52 T (7.5 N) was positioned on the right to initiate nanoculture sorting. Time-lapse imaging demonstrated that the magnetically responsive MNCs moved toward the magnetic source, displacing the non-magnetic RFP-tagged nanocultures in the process (**Figs. 5 A1-A2, Supporting Video 6)**. By the end of the sorting process, the majority of MNCs closest to the magnet were Syto 9-stained *E. coli Nissle,* as shown by tracking the trajectories of individual magnetic versus non-magnetic nanocultures (**Fig. 5 B-C)**. The magnetically actuated MNCs exhibited an average velocity of approximately 50 μm/s, a decrease attributed to the hindrance caused by interactions with non-actuated nanocultures (**Fig. 5 D)**. In contrast, the non-MNCs exhibited less controlled movement, being displaced only when pushed by the MNCs, with an average velocity of less than 20 μm/s (**Fig. 5 E)**. These results confirm that magnetic actuation enabled precise and efficient separation of the magnetically functionalized nanocultures from their non-magnetic counterparts, demonstrating a clear proof of concept for the magnetic sorting capabilities of the MNC system.

To further validate the real-world applicability of MNCs, we tested the system’s functionality in more complex, heterogeneous environments by simulating soil ecosystems (**Fig. 6**). **Fig. 6 A** provides an overview of the experimental setup, where *E. coli Nissle* containing MNCs, functionalized with 5 nm MNPs at 500 ppm, were added to the substrate. **Fig. 6 B1** shows fluorescent imaging of *E. coli Nissle* MNCs stained with Syto 9, highlighting the successful encapsulation and fluorescence tagging of the bacteria. To simulate the spatial heterogeneity encountered in natural soils, zirconia/silica beads were used as a surrogate substrate (**Fig. 6 B2).**

The experimental setup aimed to retrieve the MNCs from beneath the bead layers using a neodymium magnet placed outside the imaging area, to avoid optical interference. Our results demonstrated that within 60 min, 149 out of 151 MNCs were successfully retrieved from beneath the zirconia/silica beads, achieving a >98 % recovery rate (**Fig. 6 C1–C3, Supporting Video 7)**.

To add complexity, we used beach sand as a substrate to test whether the MNCs could sustain retrieval in a real-life soil-like system (**Fig. 6 D)**. The sand was tightly packed into a sealed hybridization chamber (**Fig. 6 E)**, and the MNCs were introduced into the chamber before being exposed to the magnetic field. Although the system was less optically clear due to the presence of sand (**Fig. 6 F)**, the MNCs were still visible aligning with the magnetic field, forming a curved trajectory corresponding to the curved shape of the magnet (**Fig. 6 G, Supporting Video 8)**. While imaging the capsules within the sand system proved more challenging than with silica beads, the MNCs were successfully retrieved within 30 min. **Fig. 6 F-G** depict images of the heterogeneous sand particles and the retrieval of MNCs from the sand environment before and after alignment. While retrieval in silica bead and sand-containing environments demonstrate real-world applicability, accurately tracking capsule motion using MATLAB was challenging due to background interference from similarly sized sand particles. To address this, we calculated the average velocity of MNCs under identical conditions but in the absence of silica beads, enabling cleaner segmentation and motion tracking. Under these conditions, the MNCs exhibited velocities ranging from approximately 100 µm/s to 140 µm/s. with an average velocity of 120 +/- 20 μm/s **(Fig. S 9, Supporting Video 9)**, though we anticipate some hindrance-induced reduction in velocity in packed environments. Nonetheless, the retrieval success demonstrates the robustness and potential of the MNC system in complex, heterogeneous environments, highlighting its future potential for environmental applications, such as microbial isolation in soil or other complex substrates.

## DISCUSSION

In this study, we developed a tunable microbial MNC system using a PDMS-based polymer matrix functionalized with MNPs. The magnetic susceptibility of the MNCs can be precisely controlled by adjusting the size and concentration of the MNPs, although nanoparticle size influences the thickness of the MNC walls. Importantly, the presence of MNPs did not impact the microbial growth of *S. aureus, P. aeruginosa, and E. coli Nissle*, demonstrating the biocompatibility of the system.

Through selection of MNP size and optimization of concentration during fabrication, we controlled the magnetic susceptibility of the MNCs, enabling velocities ranging from approximately 80 to 200 μm/s under varying field conditions. This velocity control enables the efficient separation of MNCs from non-magnetic counterparts and facilitates successful retrieval from complex environments such as silica/zirconia bead substrates and soil-like matrices. The high recovery rates (>98 %) in these heterogeneous systems highlight the robustness and real-world applicability of the MNC platform.

As advancements in environmental microbiome research continue to reveal their critical role in health and ecological systems, culture models like the one presented here are essential for improving microbial culturability. Further, this system opens new avenues for manipulating microbial dynamics and enabling targeted delivery through magnetic actuation, offering significant potential for future applications in environmental microbiology, therapeutic delivery, and microbial isolation from diverse substrates.

Our current findings highlight the system’s capabilities and point to exciting opportunities for further exploration. While we demonstrated compatibility with model pathogens and validated magnetic actuation in soil-like substrates, the system has not yet been tested with complex environmental microbiomes or under fully native soil or marine conditions. Additionally, although the platform is designed to support molecular exchange across its semi-permeable membrane, we did not directly measure small-molecule diffusion or microbial cross-communication in this study. Compared to conventional methods, further research is needed to assess microbial community dynamics, interactions, and gene expression within MNCs.

### Limitation of the study

While the MNC platform shows promise for in situ microbial assessment, its performance in complex and heterogeneous soil environments remains a key limitation. Future efforts should focus on deploying the system in field settings and validating its ability to support the growth of previously uncultured species under environmentally relevant conditions (*36*). Assessing the influence of soil properties—such as texture, coarseness, and depth (e.g., topsoil vs. subsoil; loam, sandy loam, and clay-loam)—on retrieval efficiency and magnetic responsiveness will be critical for optimizing deployment strategies (*37*). Additionally, integrating the platform with high-throughput sequencing and metabolomics could enhance its utility in characterizing the structure and function of encapsulated microbiomes. Addressing these challenges is essential to fully realize the potential of the MNC platform as a robust tool for ecological discovery, therapeutic microbiology, and precision microbiome engineering.

## MATERIALS AND METHODS

### Magnetic Nanoparticles

Iron oxide nanoparticles (MNPs) coated with oleic acid and dispersed in toluene were purchased from Cytodiagnostics Inc. (Canada). Three nominal core sizes were used: 5 nm +/- 1.5 nm, 10 nm +/- 1.5 nm, and 20 nm +/- 1.5 nm. Iron oxide MNPs were chosen for their reported ferrimagnetic properties. The oleic acid coating of the nanoparticles provided the stability and hydrophobicity required for easy mixing in PDMS blends (*38*).

### Polymer Synthesis

To synthesize polydimethylsiloxane membranes and microcapsules, commercially available vinyl (DMS–V21: *Vinyl-terminated polydimethylsiloxane*) and hydride (HMS–053: *Methylhydrosiloxane-Dimethylsiloxane Copolymer and Trimethylsiloxane Terminated*) were acquired (Gelest, Inc., USA). The hydride and vinyl were mixed in a glass vial (Grainger, USA) and further supplemented by platinum (Pt)-divinyltetramethylsiloxane (Pt in xylenes, Gelest Inc., USA) catalyst to initiate the hydrosilylation reaction and crosslinking. Previously, we determined an optimal mixing ratio of vinyl to hydride to achieve stable microcapsules for microbial encapsulation (*19–22*). Following a similar protocol with slight modification, the hydride and vinyl were mixed at a molar ratio of 0.6:1 with 1.4 +/- 0.4 ppm Pt catalyst. To generate magnetically responsive membranes with optimal stability and optical properties, iron oxide MNPs were introduced into the predefined PDMS mixture at concentrations of 0 ppm, 60 ppm, 125 ppm, 250 ppm, and 500 ppm. The composite mixtures were poured into a glass vial (Grainger, USA) and vortexed for 1 to 2 min to ensure uniform mixing before being degassed under vacuum for 5 to 10 min to remove any air bubbles. The mixtures were then used for encapsulation or cured as membranes for characterization.

### UV-Vis Characterization

To measure their optical properties, the PDMS mixtures were prepared with the 5 nm, 10 nm, and 20 nm MNPs at concentrations ranging from 0 to 500 ppm, as described above. The mixtures were introduced into spectrophotometer cuvettes (Thermofisher, USA) and briefly exposed to heat at 70 °C for 5 minutes to initiate cross-linking. The samples were incubated and cured at 37 °C for 24 h to mimic the growth conditions of model organisms used in the study. The cuvettes containing samples (path length of 1 cm and 0.4 cm) were placed in the UV-Vis spectrophotometer (Evolution 202 spectrophotometer, Thermofisher, USA) to record the percentage transmittance peaks across wavelengths ranging from 300 nm to 1000 nm.

### Contact Angle Measurements

The water contact angle of the samples was measured to determine whether the iron oxide MNPs altered the surface hydrophobicity of the PDMS membranes. Therefore, 1 μL of distilled water was applied to the membranes prepared above, using an optical tensiometer (Attention theta flex, Biolin Scientific). The sessile drop method (OneAttension) was employed to measure the respective contact angles. The experiments were repeated at least three times to ensure reproducibility.

### Superconducting Quantum Interference Device–Vibrating Sample Magnetometer Measurements (SQUID-VSM)

The composite membranes, namely PDMS with Iron Oxide MNPs, were sliced with a razor blade, weighed, and mounted with polyester tape onto a glass sample holder. The static magnetic moment versus applied magnetic field was measured at 277 K, 310 K, and 323 K from +/- 5.57 MA/m (70 kOe) in a SQUID-VSM (Quantum Design, Inc.) with an amplitude of 4 mm, at a fixed range of 100, for 2 s. A control PDMS sample without the MNPs was measured under identical conditions to permit point-by-point subtraction of the diamagnetic background. The data was normalized by the mass of the sample.

### Keyence Profilometry

The PDMS mixture containing 500 ppm MNPs was introduced into mini Petri dishes (Falcon) to yield a smooth solid sample measuring 2 mm +/- 1 mm in thickness. For nanoparticle distribution assessment within the polymer, an optical profilometer (Keyence) was employed. Using depth profiling through a 1000× magnification with digital acquisition, Z-stack images of the composite membranes were generated to elucidate the size distribution of the nanoparticles, specifically when aggregates were formed. Quantitative image-based analysis of these Z-stacks using ImageJ was used to determine the Feret diameter of these aggregates within the membranes.

### Transmission Electron Microscopy (TEM) Characterization

TEM samples were prepared by depositing 0.2 μL – 0.5 μL of MNPs as received onto ultrathin carbon-coated copper TEM grids (Ted Pella, Inc). The samples were allowed to dry overnight in a fume hood before imaging. TEM characterization was performed on a Thermo Fisher Scientific Themis G2 200 probe-corrected TEM/Scanning-TEM equipped with a Super-X EDS system. The accelerating voltage was constant at 200 kV. Under STEM imaging, a probe current of 65 pA was used. Particle size analysis was performed using a threshold-based method on high-angle annular dark-field (HAADF) STEM images. The TEM images were processed with ImageJ to quantify the particle size distribution. The total number of particles measured was 2202, 9562, and 4687 for the 5 nm, 10 nm, and 20 nm samples, respectively.

### Fabrication of Microfluidic Devices

To generate the W/O/W emulsions (the basis for the nanocultures), a glass-capillary microfluidic device was used as described previously but with slight modifications (*19–22*). Briefly, two circular capillary tubes, with inner and outer diameters of 0.58 mm and 1.03 mm (World Precision Instrument), respectively, were tapered to the desired diameters using a Sutter P-1000 Horizontal Micropipette Puller (Sutter Instrument) and an MF2 microforge (Narishige). The inner diameters of the tapered tubes for the injection of the bacteria phase and the collection of the capsules were 40 μm and 200 μm, respectively. The outside of the glass-capillary tube for the injection of the bacteria phase was hydrophobically functionalized with 1 % octadecyltrichlorosilane (OTS, Sigma-Aldrich) in toluene. This chemical treatment enhanced the wettability of the PDMS mixture outside the capillary tube and facilitated the formation of the W/O/W emulsions (*9, 19–22*). The two tapered capillary tubes were inserted into a square (inside length of a side was 1.05 mm) capillary tube with a separation of 120 μm. A transparent epoxy was used to seal the tubes, as required.

### Generation of the Magnetic Nanocultures

The microfluidic device was mounted on an inverted optical microscope (Eclipse TE300, Nikon). Then, the three fluid phases were delivered to the microfluidic device through polyethylene tubing (Scientific Commodities) attached to syringes (SGE), which were driven by positive displacement syringe pumps (Harvard Apparatus, PHD ULTRA). The drop formation was monitored with a Phantom VEO 710 high-speed camera (Vision Research) attached to the inverted microscope. The innermost fluid consisted of bacteria suspended in the culture medium, and the middle fluid of the PDMS mixture with HMS-053 and DMS-V21 (0.6:1 molar ratio) and supplemented with 1.4 +/- 0.4 ppm Pt (Gelest Inc.). The PDMS mixture was functionalized using 500 ppm magnetic nanoparticles (Cytodiagnostics Inc.). The outermost fluid consisted of a 2.5 wt.% to 5 wt.% poly-(vinyl alcohol) aqueous solution (PVA, 87% − 89% hydrolyzed, average molecular weight, Mw = 89000−98000, Sigma-Aldrich). Once the bacterial suspension was encapsulated in the magnetized PDMS mixtures, polymerization was initiated by heat treatment for 5 min at 70 °C and then incubation at 37 °C for 24 h. (This heat treatment condition was also determined to be safe for encapsulated bacteria such as *E. coli Nissle* and *P. aeruginosa* (*20, 22*). Alternatively, the background nanocultures were collected in Milli-Q water and secured in a MatTek dish before evaluating their stability for 24 h using a Zeiss Axio Imager M2 light microscope (Carl Zeiss, Germany) on a stage heated to 37 °C. A background nanoculture or microcapsules were made where the innermost fluid contained only the MilliQ sterile water, but no bacteria in culture media, as described above.

### Microorganisms and Culture Media

*E. coli Nissle* 1917 and Green Fluorescent Protein-tagged *P. aeruginosa* (PAO1) were the bacteria used in magnetic nanocultures (MNCs). The bacteria were cultured in a buffered low-molecular-weight medium referred to as ultrafiltered tryptone-yeast extract (UFTYE) broth. This UFTYE broth contained 2.5 % tryptone and 1.5 % yeast extract in water with a molecular-weight cut-off of 10 kDa (Millipore) and a pH adjusted to 7. A 25 μL aliquot of the aqueous bacterial suspension was introduced into 5 mL of UFTYE to form the aqueous suspension of bacteria in the culture media described above. Finally, the aqueous bacterial suspension encapsulated in the nanocultures was then imaged using the Zeiss Axio Imager M2 microscope for time-lapse (Carl Zeiss, Germany) or Leica STELLARIS 5 for confocal end-point imaging (Leica, USA).

### Scanning Electron Microscopy (SEM) Characterization

SEM was employed to analyze the architecture of the nanocultures, both with and without MNPs. The fully cured concentrated nanoculture solution was transferred to a 15 mL conical tube containing 10 % glycerol in sterile MilliQ water. The conical tube underwent centrifugation at 6000 RPM for 10 min, causing the nanocultures to settle to the bottom due to density differences. Subsequently, the nanocultures were transferred into a small glass vial and subjected to a 48 h lyophilization process. The lyophilized materials were then mounted on conductive carbon adhesive tape and sputter-coated with a gold-palladium alloy for electrical conductivity before imaging, using the Zeiss SIGMA VP electron microscope at an accelerating voltage of 3 kV.

### Magnetophoresis Experimental Setup

To evaluate their magnetophoretic properties, 100 μL to 200 μL of concentrated and fully polymerized nanocultures were added to a hybridization collection chamber (ThermoFisher, USA). The chambers were prefilled with sterile UFTYE culture media or MilliQ water for the background nanoculture. The medium in the collection chambers was maintained to be similar to the core of the nanocultures to prevent osmotic changes and subsequent magnetophoretic disturbances. The hybridization chamber was mounted on an inverted optical microscope (Eclipse TE300, Nikon) to visualize the samples (Eclipse TE300, Nikon). Neodymium iron boron magnet (nickel-coated, 0.635 cm thick, 2.54 cm OD, McMaster-Carr) with surface magnetic induction fields of 1.48 T was used to induce field-dependent magnetophoresis. When cell sorting with magnetophoresis was performed, the nanocultures with actuation with MNPs were stained with Syto 9, selected due to its ability to readily diffuse through the nanoculture and permeate bacterial cell membranes, staining all nucleic acids green, thus making them visible for fluorescent microscopy (*9, 21*).

### Tracking Motion of MNCs

The motion of the MNCs was recorded using a high-speed camera at a frame rate of 10,000 fps with a 4× objective (Phantom VEO 710L from Vision Research). The motion of the MNCs with bacteria was recorded using a fluorescent microscope with a monochrome camera at a frame rate of 2-3 fps (Axiocam 702 in burst mode from Carl Zeiss). Custom MATLAB code was used to track the trajectories of MNCs through the recorded image sequences in the following manner: First, the MNCs were identified in recorded images by employing Otsu thresholding (*39*) to binarize images and separate regions of MNCs from the background. Morphological transformation was then used to determine the center of each MNC (*40*). The centers of MNCs were tracked over time relative to the magnet using the Crocker–Grier multiple-particle tracking algorithm (*41*). The code then identified the unique particle IDs, and for each particle with sufficient data points, pixel coordinates were converted to microns, and the time in seconds was calculated. Differences in *x* and *y* positions between consecutive frames were computed, enabling the calculation of velocities in both directions (no significant differences in y positions were identified). Velocity data was aggregated, and average velocities at different positions along the channel length and width were calculated, using median and standard deviation filtering to remove outliers.

### Statistical Analysis

GraphPad Prism Software (V8) was used to conduct the t-test, one-way ANOVA, and two-way ANOVA to establish the significance of the testing conditions. Differences of p < 0.05 were considered statistically significant. The following notations “ns”, *, **, ***, and **** describe the statistical difference with p values corresponding to p > 0.05, p < 0.05, p < 0.01, p < 0.001, and p < 0.0001, respectively.

## Acknowledgments

We gratefully acknowledge the independent reviewers at NIST for their insightful evaluations and constructive feedback as part of the internal review process. Electron microscopy was conducted at the Nanoscale Fabrication and Characterization Facility (NFCF) within the Petersen Institute of NanoScience and Engineering (PINSE) at the University of Pittsburgh. This work also benefited from the Environmental TEM Catalysis Consortium (ECC), supported by the University of Pittsburgh and Hitachi High Technologies. We thank Dr. Simon C. Watkins and the Center for Biologic Imaging for providing access to scanning electron microscopy resources.

## Authors Contributions

Conceptualization: THRN, MFH, HU

Methodology: HU, THRN, SH, CLD, MM

Investigation: HU, THRN, SH

Visualization: HU, THRN

Supervision: THRN, MM, CLD

Writing—original draft: HU, MM, SH, CLD, MFH, THRN

Writing—review & editing: HU, MM, SH, CLD, MFH, THRN

## Data and materials availability

All data that support the findings of this study are available in the main text or the supplementary materials. Additional questions and all other correspondence should be addressed to.

## Disclaimer

We identify certain commercial equipment, instruments, or materials in this article to specify adequately the experimental procedure. In no case does such identification imply recommendation or endorsement by the National Institute of Standards and Technology, nor does it imply that the materials or equipment identified are necessarily the best available for the purpose.

