## Supporting Information for "Magnetically Responsive Nanocultures for Direct Microbial Assessment in Soil Environments"

Science Advances
AAAS

### Supplementary Materials for

#### Magnetically Responsive Nanocultures for Direct Microbial Assessment in Soil Environments

Huda Usman *et al.*

**This PDF file includes:**

Supplementary Text
Figs. S1 to S9
Tables S1
Movies S1 to S8

**Other Supplementary Materials for this manuscript include the following:**
Movies S1 to S8

**Supplementary Text**

In this supplementary information, we have included a summary of statistics of size distributions, diffraction patterns, and energy-dispersive X-ray spectroscopy spectra of the iron oxide nanoparticles (MNPs) referred to in the main text. The MNPs embedded polymer was also characterized for its hydrophobicity, which is also discussed. Further, we have elucidated the effect of MNP exposure for 0 ppm, 60 ppm, 125 ppm, 250 ppm, and 500 ppm on the growth dynamics of cells by comparing the lag time, μmax, and growth potential versus the untreated control. Representative videos for the stability and magnetophoresis of the microcapsules made with 5 nm MNPs are also included. NCs generated using the 500 ppm 5 nm MNPs containing *E. coli Nissle* video are also included, and the average velocity of the MNCs is determined. For readability, the following notations "ns", *, **, ***, and **** describe the statistical difference with p values corresponding to $p > 0.05$, $p < 0.05$, $p < 0.01$, $p < 0.001$, and $p < 0.0001$, respectively and the differences of $p < 0.05$ were considered statistically significant.

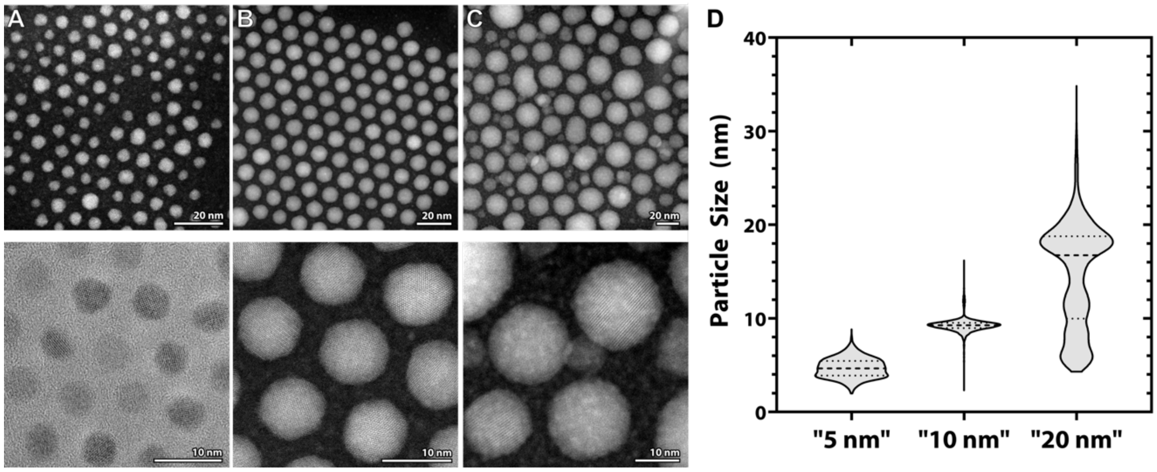


**Fig. S1. Characterization of the MNP size distribution using TEM.** HAADF-STEM overview images showing the range of sizes and morphology of the (**A**) 5 nm, (**B**) 10 nm, and (**C**) 20 nm samples, respectively (Top Panel). HR-TEM and HAADF-STEM close-up images of the particles, showing their crystallinity (Bottom Panel); (**D**) Violin plot of measurement particle size. For each distribution, the dotted line marks the median and the dashed lines mark the 25 % and 75 % interquartile ranges (IQR).

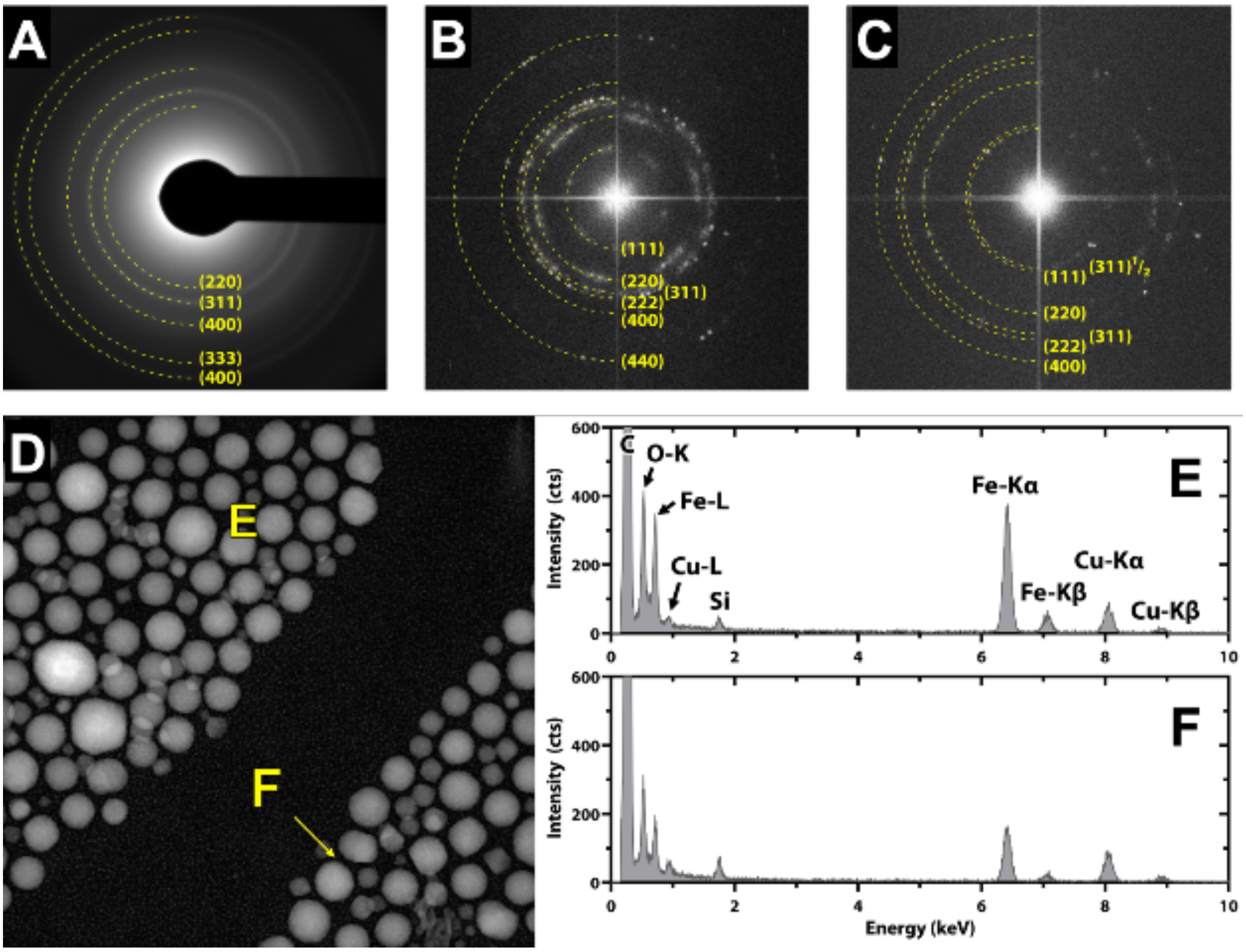


**Fig. S2. X-ray diffraction (XRD) and energy-dispersive X-ray spectroscopy (EDS) measurements of the MNPs.** (**A** – **C**) Diffraction patterns from samples of the 5 nm, 10 nm, and 20 nm magnetic MNPs. (**D**) HAADF-STEM micrograph from the 20 nm MNP-PDMS sample. (**E, F**) Labeled energy-dispersive X-ray spectroscopy spectra from a (**E**) large round and (**F**) small, faceted nanoparticle, as indicated in (**D**).

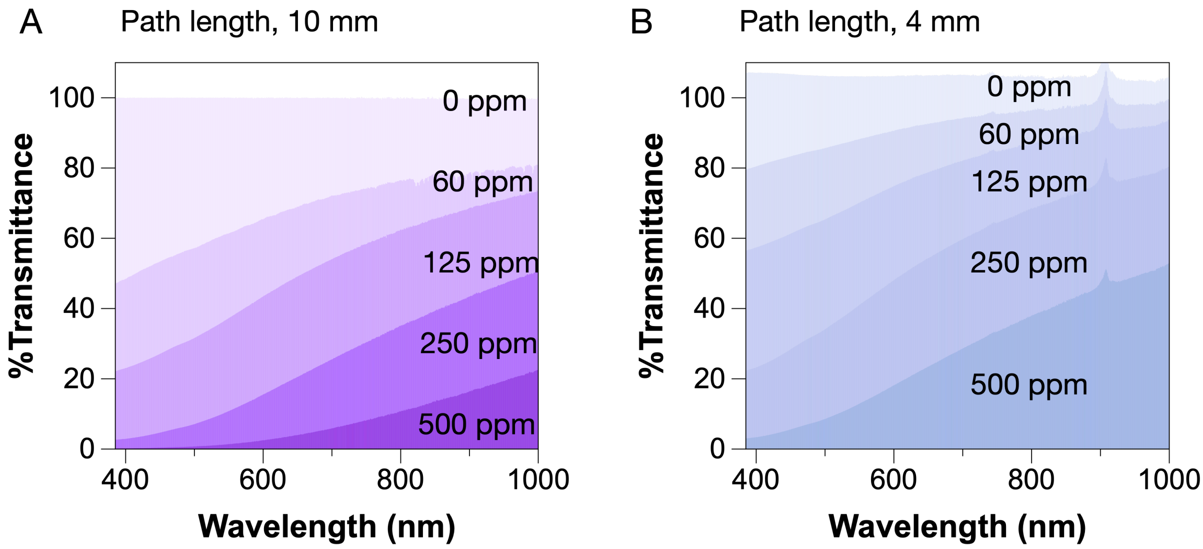


**Fig. S3. Transmittance as a function of wavelength for polymer blends containing 5 nm MNPs at varying concentrations and path lengths**. (**A**) The transmittance of polymer blends containing 5 nm MNPs was measured across the visible spectrum (385 nm – 750 nm) for the path length of 10 mm at different MNP concentrations (60 ppm, 125 ppm, 250 ppm, and 500 ppm), (**B**) The transmittance of polymer blends containing 5 nm MNPs was measured across the visible spectrum (385 nm – 750 nm) for for the path length of 4 mm at varying MNP concentrations.

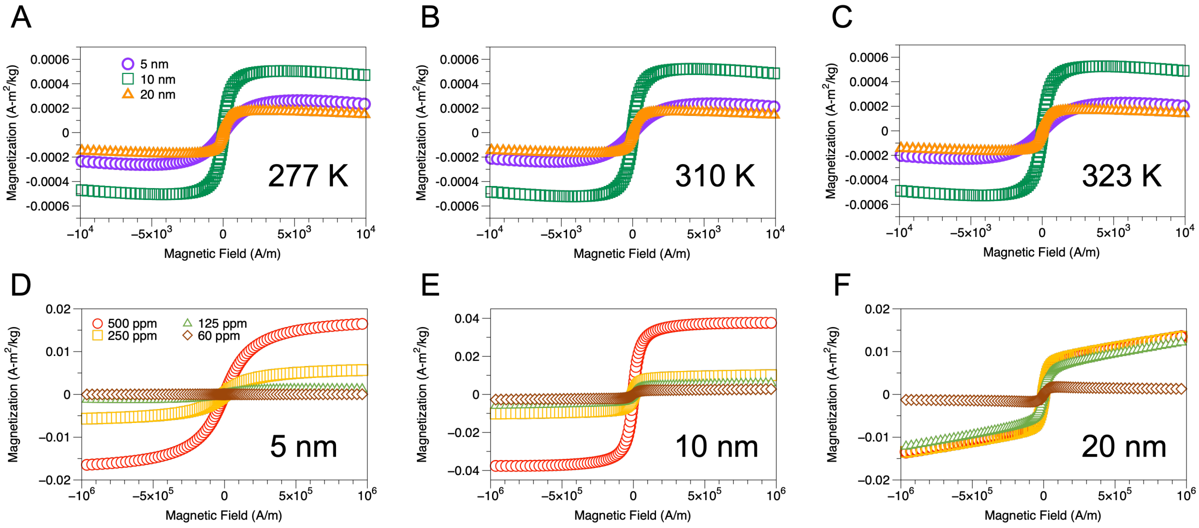


**Fig. S4. Characterization of Magnetic Properties of the Polymer Blends at Varying Concentrations and Temperature** (**A-C**) Hysteresis curves of polymer-MNP blend with 500 ppm concentration at additional temperatures (277 K, 330 K, and 323 K) illustrate the influence of temperature on magnetic behavior, with temperature variations affecting both the coercivity and magnetic saturation, (**D**) Magnetization curves for polymer blends containing 5 nm MNPs measured at 310 K across different concentrations (60 ppm, 125 ppm, 250 ppm, and 500 ppm) show a gradual increase in magnetic response with concentration, (**E**) Magnetization curves for blends with 10 nm MNPs at 310 K indicate a similar concentration-dependent increase in magnetization, with higher saturation observed than with 5 nm particles, (**F**) The magnetic response of 20 nm MNP-embedded blends at 310 K further emphasizes the effect of particle size on saturation magnetization.

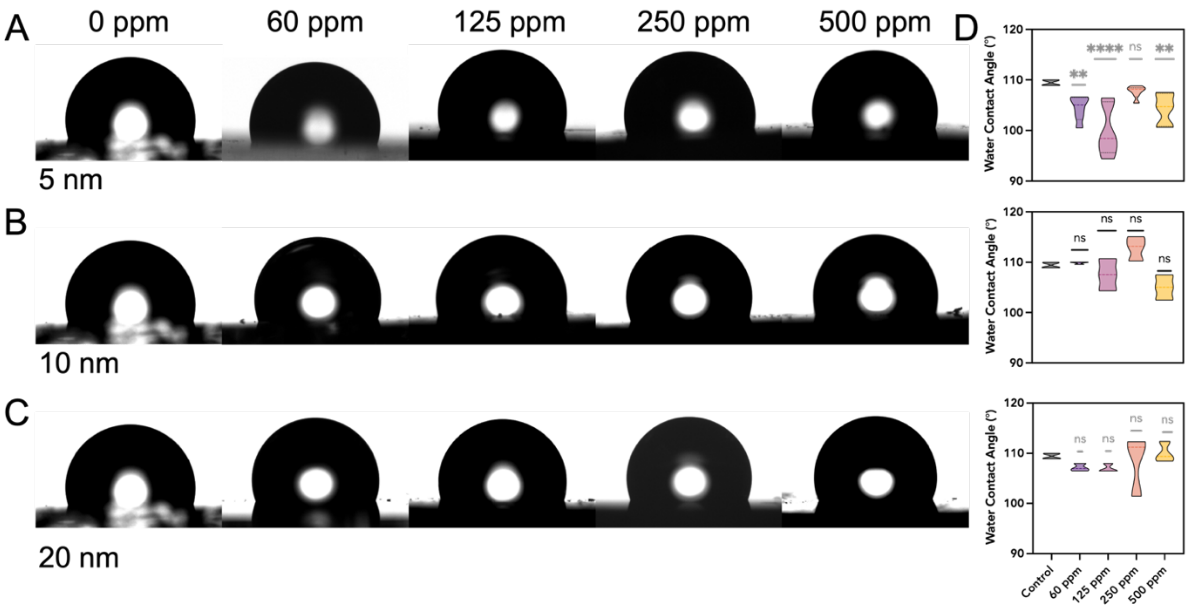


**Fig. S5.** Contact angle analysis of PDMS membranes functionalized with magnetic nanoparticles. (**A–C**). Representative images of 1–2 µL water droplets on PDMS membranes functionalized with varying concentrations (0 ppm, 60 ppm, 125 ppm, 250, and 500 ppm) of (**A**) 5 nm, (**B**)10 nm, and (**C**)20 nm MNPs, respectively. The 0 ppm membrane serves as the non-functionalized control. (**D**) Quantified absolute contact angles measured for each condition. A two-way ANOVA revealed statistically significant effects of both MNP size and concentration ($p < 0.0001$), as well as a significant interaction between the two ($p = 0.0039$). Post hoc Dunnett's tests showed that only 5 nm MNPs at 60 ppm, 125 ppm, and 500 ppm resulted in significantly lower contact angles compared to the control, indicating increased hydrophilicity at those concentrations.

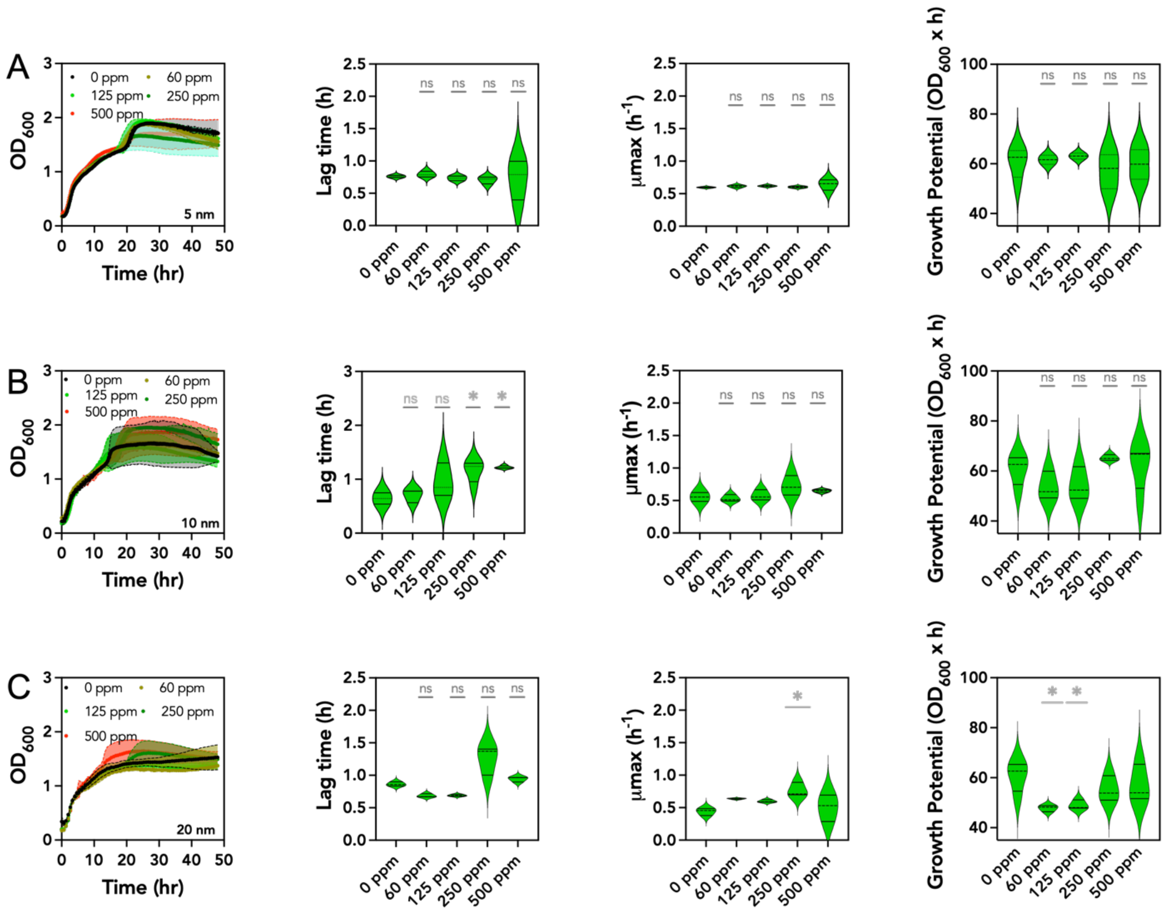


**Fig. S6.** Cell susceptibility when exposed to varying 5 nm, 10 nm, and 20 nm MNP concentrations. (**A-C**) *Staphylococcus aureus* was exposed to MNP concentrations of 0 ppm, 60 ppm, 125 ppm, 250 ppm, and 500 ppm. Growth curves were constructed by taking OD measurements every 10 min over 48 h. The growth curves were analyzed using R Studio (a statistical computing environment). Each curve was fitted into a logarithmic equation, and the lag time, the μmax, and the area under the curve were integrated. The values were compared to estimate statistical differences.

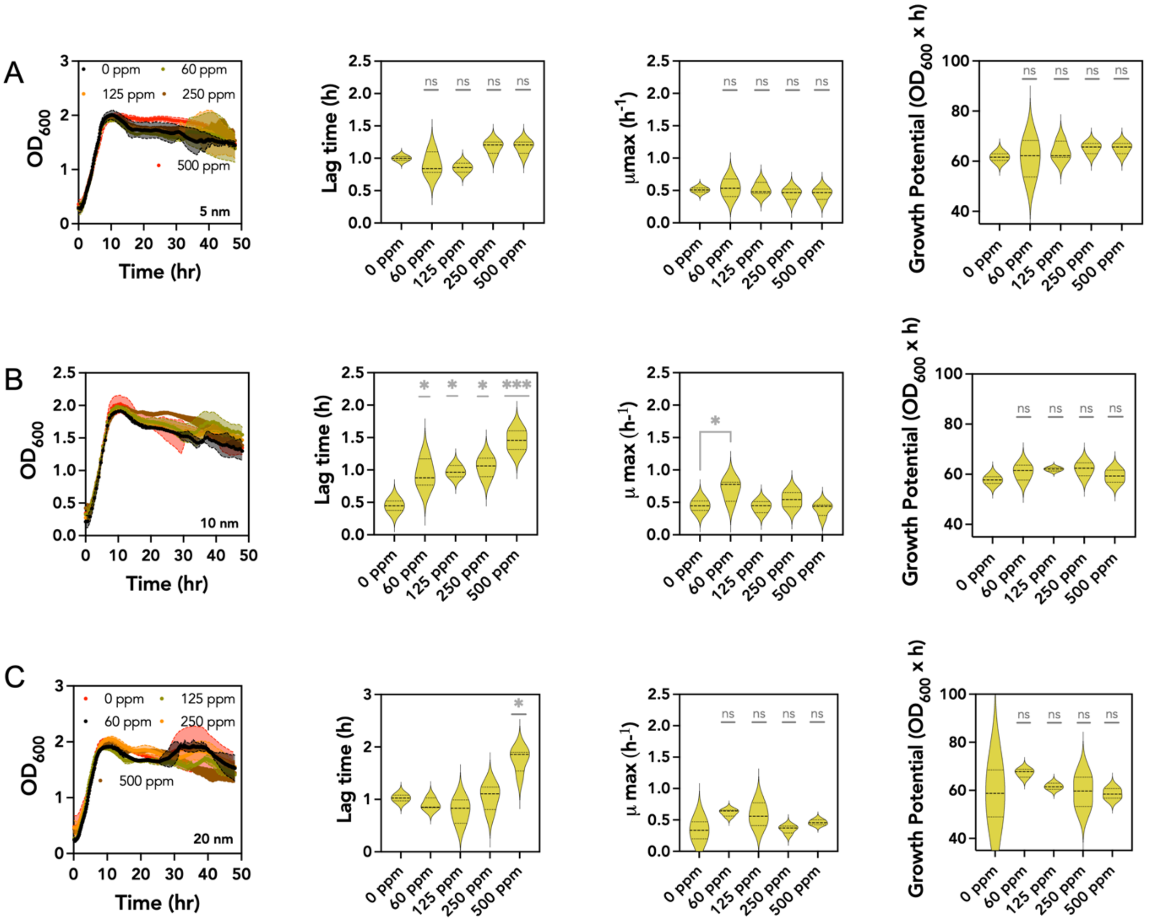

**Fig. S7.** Cell susceptibility when exposed to varying 5 nm, 10 nm, and 20 nm MNPs concentrations.

(**A-C**) *Pseudomonas aeruginosa* was exposed to MNP concentrations of 0 ppm, 60 ppm, 125 ppm, 250 ppm, and 500 ppm. Growth curves were constructed by taking OD measurements every 10 min over 48 h. The growth curves were analyzed at R Studio (a statistical computing environment). Each curve was fitted into a logarithmic equation, and the lag time, the μmax, and the area under the curve were integrated. The values were compared to estimate statistical differences.

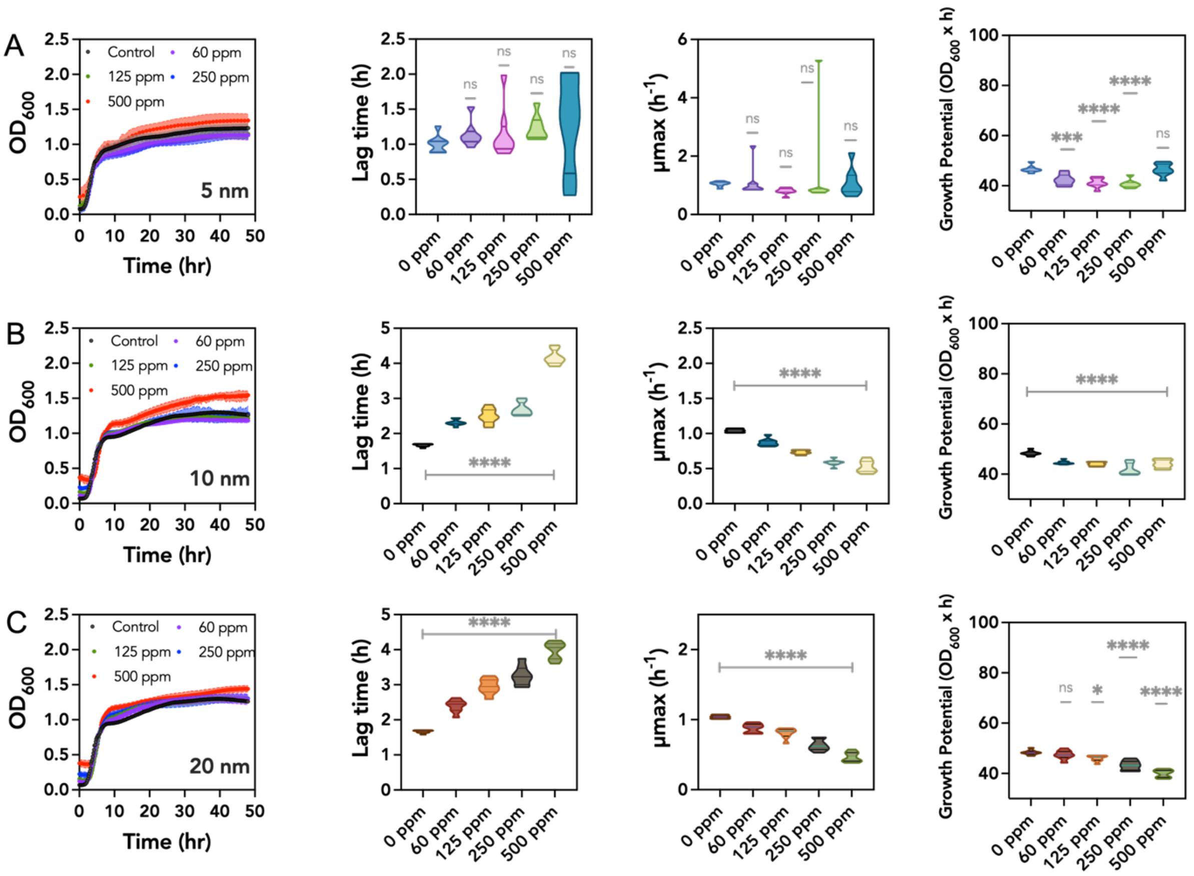


**Fig. S8.** Cell susceptibility when exposed to varying 5 nm, 10 nm, and 20 nm MNPs concentrations. (**A-C**) *Escherichia coli Nissle* was exposed to MNP concentrations of 0 ppm, 60 ppm, 125 ppm, 250 ppm, and 500 ppm. Growth curves were constructed by taking OD measurements every 10 min over 48 h. The growth curves were analyzed at R Studio (a statistical computing environment). Each curve was fitted into a logarithmic equation, and the lag time, the μmax, and the area under the curve were integrated. The values were compared to estimate statistical differences.

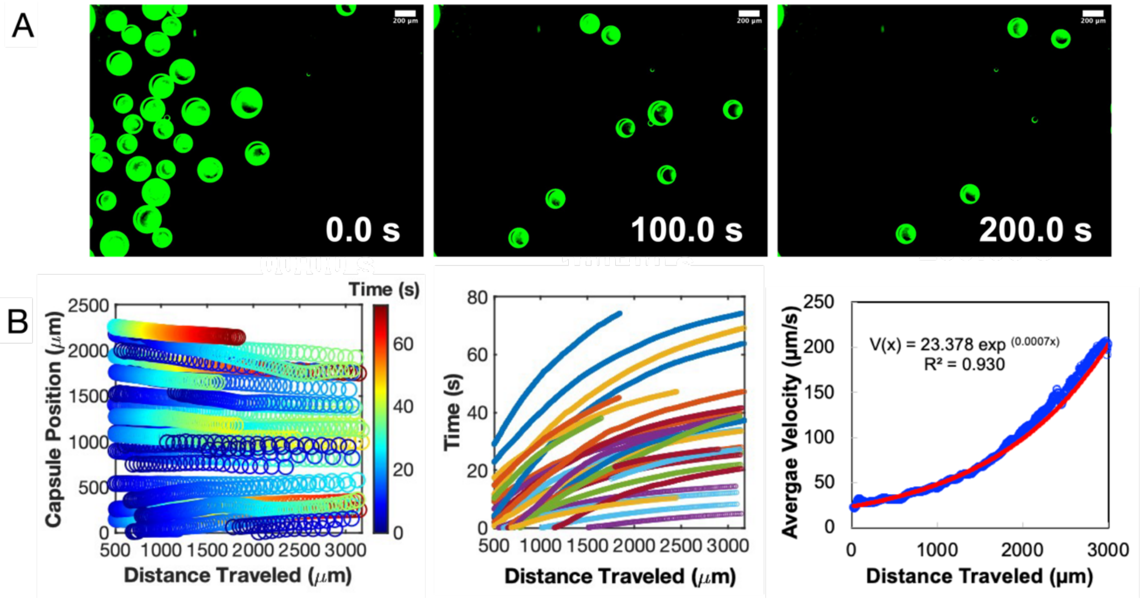


**Fig. S9.** Magnetic actuation of *E. coli Nissle* nanocultures without the presence of silica beads: (**A**) Movie captured from a brightfield camera showing the ability of the nanocultures to move toward the magnet. Scale bar: 200 μm. (**B**) The plots show MATLAB-rendered trajectories of the nanocultures as a function of their distance from the magnet and their average velocity gradient (~120 μm/sec).

| | Iron oxide nanoparticles | | |
|---|---|---|---|
| | **5 nm** | **10 nm** | **20 nm** |
| **# of MNPs Measured** | 2202 | 9562 | 3220 |
| **Mean (nm)** | 4.7 | 9.3 | 14.8 |
| **Std. Dev. (nm)** | 1.0 | 0.9 | 5.5 |
| **25 % IQR (nm)** | 3.9 | 9.0 | 10.0 |
| **Median (nm)** | 4.7 | 9.3 | 16.8 |
| **75 % IQR (nm)** | 5.5 | 9.5 | 18.8 |

**Table S1.** Particle size summary statistics of the distributions are plotted in Fig. S1, D.

**Supporting Movie 1**. **The stability of the microcapsules over 24 h.**

The stability of the bulk microcapsules containing water functionalized with 500 ppm 5 MNPs (MOV).

**Supporting Movie 2**. **The magnetic nanocultures undergo magnetophoresis.**

The magnetic behavior of the bulk microcapsules containing water functionalized with 500 ppm 5 MNPs (MOV).

**Supporting Movie 3**. **Growth dynamics of *E. coli Nissle* in magnetic nanocultures.**

Growth of *E. coli Nissle* cells in magnetic nanocultures recorded for 20 h is shown. A confluent bacterial growth is achieved in nanocultures functionalized with MNPs, suggesting a robust and inert nanoculture. Scale bar: 50 μm (MOV).

**Supporting Movie 4**. **Magnetic actuation of magnetic microcapsules.**

Magnetophoretic behavior of the microcapsules and nanocultures in the presence of an applied external magnetic field (MOV).

**Supporting Movie 5**. **Magnetic actuation of magnetic nanocultures.**

Magnetophoretic behavior of the microcapsules and nanocultures in an applied external magnetic field (MOV).

**Supporting Movie 6. Microbial communities sorting using magnetic actuation.**

Magnetic nanocultures containing distinct species sorting via magnetic actuation (MOV).

**Supporting Movie 7. Retrieving magnetic nanocultures containing *E. coli Nissle* in the presence of silica beads.**

Magnetic nanocultures containing *E. coli Nissle* in the presence of silica beads (MOV).

**Supporting Movie 8. Real-world applicability of magnetic nanocultures.**

Proof of concept of the applicability of our MNCs in real-world environments, particularly soil ecosystems (MOV).

**Supporting Movie 9. Retrieving magnetic nanocultures containing *E. coli Nissle* in the absence of silica beads for quantification.**

Magnetic nanocultures containing *E. coli Nissle* in the absence of silica beads for velocity calculations (MOV).
